# Molecular states underlying neuronal cell type development and plasticity in the whisker cortex

**DOI:** 10.1101/2024.10.07.617106

**Authors:** Salwan Butrus, Hannah R. Monday, Christopher J. Yoo, Daniel E. Feldman, Karthik Shekhar

**Affiliations:** Department of Chemical and Biomolecular Engineering, University of California, Berkeley, Berkeley, CA 94720, USA; Department of Neuroscience, University of California, Berkeley, Berkeley, CA 94720, USA; Helen Wills Neuroscience Institute, University of California, Berkeley, Berkeley, CA 94720, USA; Center for Computational Biology; Vision Sciences and Optometry; University of California, Berkeley, Berkeley, CA 94720, USA

## Abstract

Mouse whisker somatosensory cortex (wS1) is a major model system to study the experience-dependent plasticity of cortical neuron physiology, morphology, and sensory coding. However, the role of sensory experience in regulating neuronal cell type development and gene expression in wS1 remains poorly understood. We assembled and annotated a transcriptomic atlas of wS1 during postnatal development comprising 45 molecularly distinct neuronal types that can be grouped into eight excitatory and four inhibitory neuron subclasses. Using this atlas, we examined the influence of whisker experience from postnatal day (P) 12, the onset of active whisking, to P22, on the maturation of molecularly distinct cell types. During this developmental period, when whisker experience was normal, ∼250 genes were regulated in a neuronal subclass-specific fashion. At the resolution of neuronal types, we found that only the composition of layer (L) 2/3 glutamatergic neuronal types, but not other neuronal types, changed substantially between P12 and P22. These compositional changes resemble those observed previously in the primary visual cortex (V1), and the temporal gene expression changes were also highly conserved between the two regions. In contrast to V1, however, cell type maturation in wS1 is not substantially dependent on sensory experience, as 10-day full-face whisker deprivation did not influence the transcriptomic identity and composition of L2/3 neuronal types. A one-day competitive whisker deprivation protocol also did not affect cell type identity but induced moderate changes in plasticity-related gene expression. Thus, developmental maturation of cell types is similar in V1 and wS1, but sensory deprivation minimally affects cell type development in wS1.

**Highlights:**

- A single-nucleus transcriptomic atlas of the whisker somatosensory cortex (wS1) during early postnatal development
- Different neuronal subclasses in wS1 show distinct developmental gene expression changes
- The composition of L2/3 glutamatergic neurons changes between the second and the third postnatal week
- Developmental gene expression and cell type changes are conserved between wS1 and the primary visual cortex (V1)
- Unlike V1, these changes are not affected by prolonged sensory deprivation
- Brief whisker deprivation induces subclass-specific activity-dependent gene expression in a whisker column-specific fashion

## INTRODUCTION

Neural circuits and function in the neocortex develop in a two-step sequence. Intrinsic genetic programs specify diverse cell types and organize them into an embryonic connectivity map. Subsequently, sensory experience extensively refines this circuitry through activity-dependent mechanisms (1). The brain peaks in such experience-dependent plasticity during developmental stages known as critical periods (CPs) (2,3). These experience-dependent changes modulate individual neuronal features such as morphology and synaptic connectivity and alter network properties, including sensory coding and population activity dynamics (4–6). While sensory experience during critical periods influences the development of some neural cell populations, how these effects differ among the thousands of cell types that populate the mammalian cortex remains poorly explored (7–9).

Recent work in mouse primary visual cortex (V1) suggested that the maturation of glutamatergic neuronal types within the upper cortical layers (L2/3/4), but not lower-layer glutamatergic neurons or inhibitory neuronal types, is vision-dependent (10,11). In response to visual deprivation, the transcriptomic profiles, spatial gene expression patterns, and functional tuning of L2/3 glutamatergic neurons were altered. Whether such selective influence of experience on cell type maturation is generally conserved across neocortical areas has not been studied. Given well-described deficits in experience-dependent forms of plasticity in neurodevelopmental disorders like autism, understanding the influence of experience in normal brain development is particularly important.

To address this gap, we studied the transcriptomic maturation of cell types in the mouse whisker primary somatosensory cortex (wS1; **Fig 1A**) during normal postnatal development and during deprived whisker sensory experience. wS1 processes tactile (touch) information from the facial whiskers. This is possible because wS1 contains a somatotopic map of the whisker pad so that sensory manipulation (plucking or activation) of specific whiskers drives plasticity in the cortical columns corresponding to those whiskers. The somatotopic organization provides strong technical advantages for investigating mechanisms of plasticity and has made wS1 a workhorse for studying morphological and physiological experience-dependent plasticity (5,12,13). Cortical circuit development and critical periods are also well described in wS1 (**Fig 1B**). Thalamocortical axons arrive in L4 at postnatal day (P) 1-2, segregate into whisker-specific clusters at P3, and drive patterning of postsynaptic L4 neurons into modules (termed barrels) in L4. Barrel pattern development is partly driven by neural activity during this early period (P0-3). After this age, the anatomical barrel pattern remains stable, but alterations in whisker use drive physiological plasticity in wS1 that is maximal during two overlapping critical periods (CP1 and CP2). CP1 and CP2 initiate at the onset of active whisking (at P12) and coincide with peak synaptogenesis in wS1. From P12-14, whisker deprivation (WD) disrupts receptive field structure (14,15) and spine motility (16) in L2/3 pyramidal (PYR) cells (CP1). From P12-16, removing all but one whisker strengthens L4-L2/3 and L2/3-L2/3 synapses to enhance the representation of the spared whisker (CP2) (17). Brief 1-day whisker deprivation (WD) also drives physiological plasticity in parvalbumin-positive (PV) interneurons in L2/3, which acts to homeostatically preserve whisker-evoked firing rates in PYR cells, a form of plasticity that is robust at P21 (18). It is unknown whether these physiological changes are associated with changes in gene expression.

**Figure 1.**
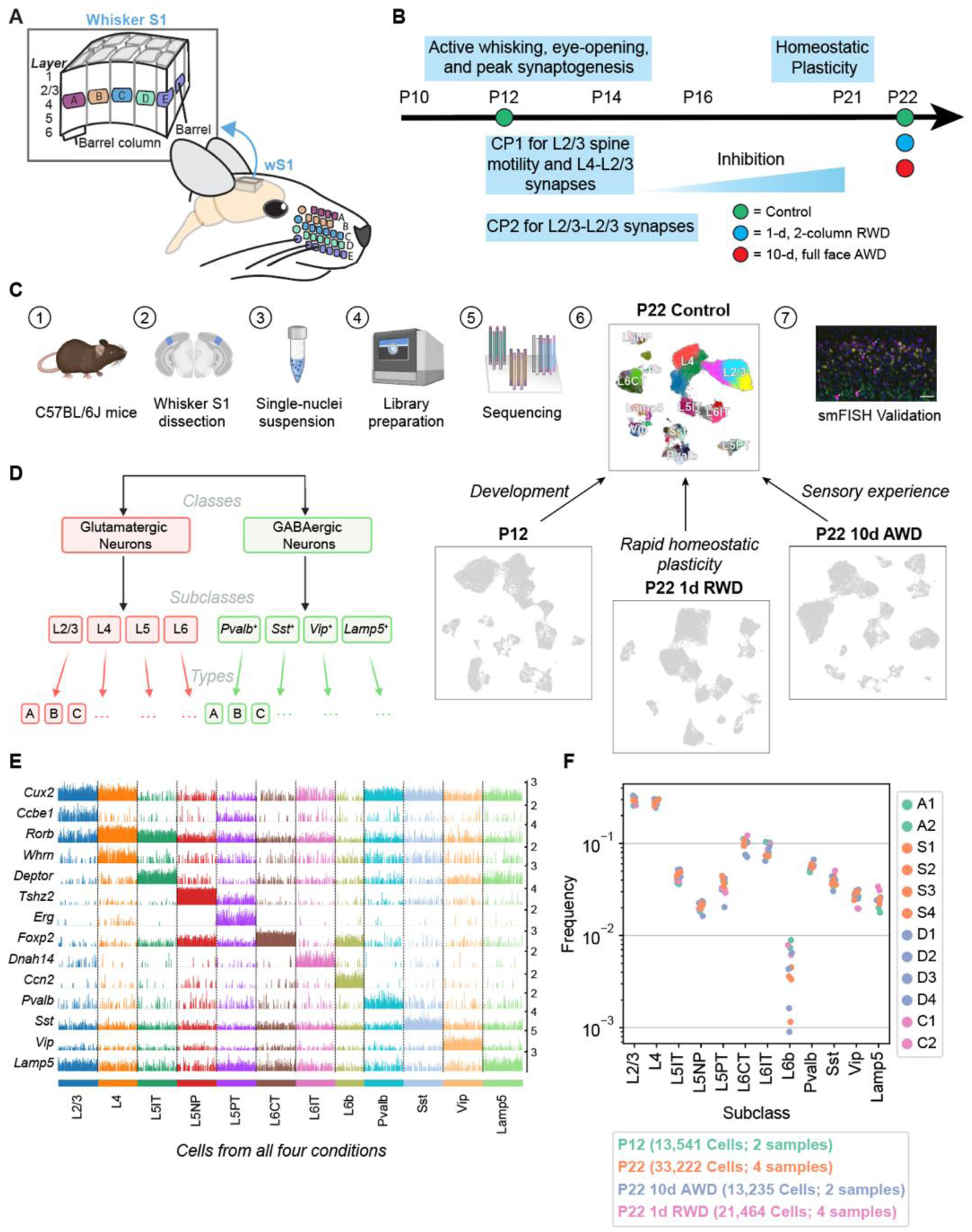
snRNA-seq atlas of the juvenile mouse primary whisker somatosensory cortex (wS1). (**A**) Schematic of the mouse whisker somatosensory system, including the facial whisker pad and the whisker somatosensory cortex (wS1). wS1 contains a somatotopic map of the whisker pad in which individual whiskers are represented by neural activity within barrel columns of the cortex. (**B**) Experimental design and developmental timeline for snRNA-seq profiling of one reference (control) dataset at P22 and three experimental conditions: an earlier time during development (P12) and following two different whisker deprivation paradigms at P22. RWD, row-whisker deprivation. AWD, all-whisker deprivation. (**C**) General experimental and computational workflow for snRNA-seq profiling and subsequent confirmatory studies. (**D**) Representation of cortical neuron diversity explored in this study highlighting the three taxonomic levels: classes, subclasses, and types. (**E**) Tracksplot showing marker genes (rows) for each neuronal subclass (columns). Data was aggregated from 81,456 nuclei across all four conditions and each subclass was subsampled to the size of the smallest subclass for plotting purposes. (**F**) Relative frequencies of neuronal subclasses are highly consistent across biological replicates and experimental conditions. The highest variance is seen for L6b glutamatergic neurons, whose frequency ranges from 0.1% to 1% of all neurons.

To study the relative contribution of intrinsic development and whisker experience to the establishment of wS1 cell types during early postnatal development, we combined single-nucleus RNA-sequencing (snRNA-seq), computational analyses, sensory perturbations, and fluorescence *in situ* hybridization (FISH) (**Fig 1C**). We first constructed transcriptomic atlases of wS1 at P12 and P22, which span from the onset of active whisking through CP1 and CP2. Within each atlas, single-nucleus transcriptomes were classified into a hierarchy comprising neuronal classes, subclasses, and types (**Fig 1D**). A comparison of gene expression between P12 and P22 identified 250 genes that were predominantly up- or down-regulated temporally in a subclass-specific fashion. We used FISH to validate some of the observed temporally regulated genes. These changes overlapped significantly with developmental changes observed in V1 during a similar period among corresponding neuronal groups (10), suggesting that postnatal maturation programs may be conserved across cortical regions. At the resolution of neuronal cell types, we found the composition of lower-layer glutamatergic and GABAergic neuronal types to be highly concordant between P12 and P22. Similar to V1, only glutamatergic neuronal types in L2/3 underwent a significant compositional shift, suggesting they mature postnatally. Surprisingly, full-face whisker deprivation from P12 to P22 had minimal effect on gene expression programs and the development of cell type composition. This contrasts with V1, where dark-rearing imparted significant gene expression and compositional changes in L2/3 (10,11). Brief (1-d) whisker deprivation of a subset of whiskers in adolescent (P21) mice, a manipulation known to drive competitive whisker map plasticity, led to whisker column-specific changes in the expression of immediate-early genes (IEGs) in L2/3, demonstrating plasticity-related regulation of gene expression. Altogether, our results suggest that wS1 and V1 develop similar cell types with a similar developmental timeline and exhibit activity-dependent gene expression changes but with very different effects of postnatal sensory experience on cell type development.

## RESULTS

### A single-nucleus transcriptomic atlas of the developing mouse whisker cortex

To begin, we used droplet-based snRNA-seq to establish a reference atlas of wS1 neurons at P22 in mice with normal whisker experience. By P22, all established CPs are complete (**Figs 1B,C**). Next, to evaluate the influence of natural development and whisker experience on neuronal transcriptomic profiles, we used the same experimental approach to profile wS1 in three scenarios to compare with the reference atlas (**Fig 1B**). First, to identify temporally regulated genes and study cell type maturation from the onset of active whisking, we profiled wS1 from whisker-intact mice at P12, which coincides with peak synaptogenesis and the onset of CP1. We performed two additional experiments to study the role of whisker deprivation in regulating cell type-specific gene expression in wS1. In one experiment, we bilaterally deprived mice by plucking all whiskers from P12 to P22 to determine if WD from the onset of active whisking affects the composition of cell types and/or their gene expression programs. In the second, we plucked the B- and D-row whiskers for 1 day, from P21 to P22. This sensory manipulation induces plasticity of excitatory and inhibitory circuits within deprived whisker columns (18,19). The goal of this experiment was to test whether a brief WD caused cell type-specific transcriptomic changes that could explain this physiological plasticity.

Data from each experiment consisted of 2-4 snRNA-seq biological replicates, each derived from cells collected from 2-3 mice (**Methods**). The resulting gene expression matrices were filtered to remove low-quality cells and cell-doublets (20), cells from non-neuronal populations, cells with a high proportion of mitochondrial transcripts (>1%), and cells that mapped poorly to other cortical datasets (**Methods**). In total, we obtained 81,462 high-quality nuclear transcriptomes corresponding to neurons across the four conditions profiled (**Figs 1C, S1A-C**). We then used standard dimensionality reduction and clustering approaches to hierarchically taxonomize wS1 neurons into 2 classes, 12 subclasses, and 53 molecularly distinct neuronal types. Annotations were performed by leveraging known markers, natural clustering, and supervised mapping to established cortical datasets (**Figs 1D-E, Methods)**. The relative frequencies of the neuronal subclasses, which spanned two orders of magnitude, were highly consistent across the experimental conditions (**Fig 1F**).

### Developmental gene expression changes are subclass-specific

Next, we sought to characterize the patterns of developmental gene expression changes across the neuronal subclasses. We computed two scores for each gene: a subclass variability (SV) score, based on its maximum expression fold change among the subclasses, and a temporal differential expression (tDE) score, based on its maximum fold change between P12 and P22 among the subclasses. A total of ∼4000 genes, each expressed in >40% of cells in at least one time point and at least one subclass, were included in this analysis (**Table S1**). Based on their scores (fold-change (FC) >2 along each axis), genes were stratified into four quadrants Q1-Q4 (**Fig 2A**, **Methods**). Glutamatergic and GABAergic subclasses showed a similar proportion of genes (∼70-80%) with low SV and tDE scores (Q3 in **Fig 2B**). However, compared to GABAergic subclasses, glutamatergic subclasses were >3x more enriched (5.3% vs 1.5%) in genes with high subclass-variability and high tDE (Q1) and 1.5x more enriched (4.5% vs 3%) in genes with low subclass-variability and high tDE (Q2). Together, these results indicate that glutamatergic neurons undergo greater transcriptional changes during this period of postnatal maturation.

**Figure 2.**
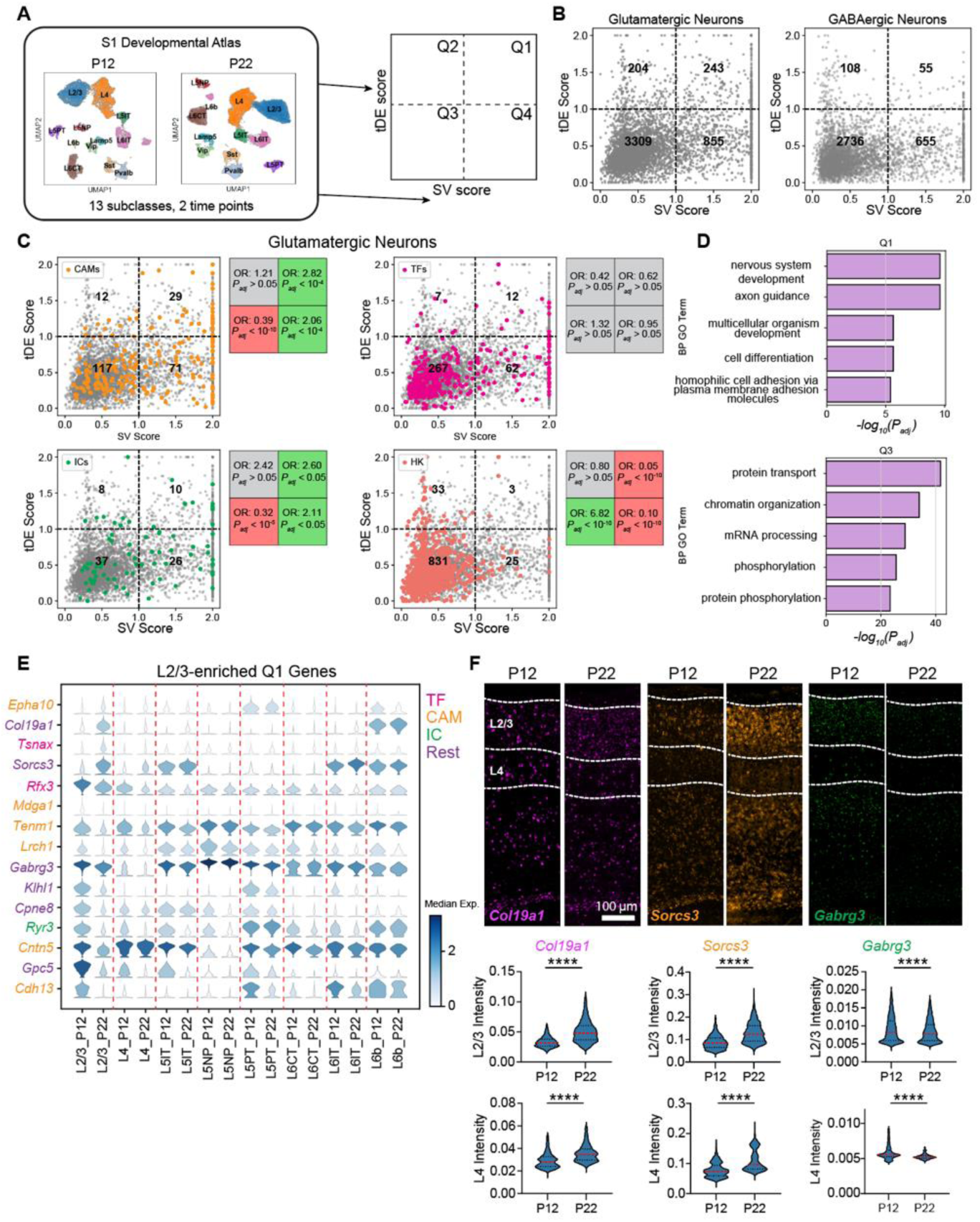
Gene expression changes between P12 and P22 are enriched in neurodevelopmental processes. (**A**) Overview of analyses to classify genes based on subclass variability and temporal differential expression. (**B**) Scatter plot of subclass variability (SV) scores and temporal differential expression (tDE) scores of genes in glutamatergic (*left*) and GABAergic neurons (*right*). Scores along each axis are capped at the value of 2. (**C**) Same as B for glutamatergic neurons with four gene categories highlighted. Subpanels (clockwise, starting from top left) correspond to cell adhesion molecules (CAMs), transcription factors (TFs), housekeeping genes (HKs), and ion channels (ICs). Boxes on the right of each panel list the odds ratio (OR) and adjusted p-value (*P_adj_)* for the enrichment of the corresponding gene category in each quadrant (Fisher’s exact test). Grey values indicate neither enrichment nor depletion, while red and green indicate depletion and enrichment, respectively (see **Methods** for details). (**D**) Q1 is enriched in gene ontology (GO) programs associated with neurodevelopment, while Q3 is enriched in general housekeeping processes. (**E**) Stacked violin plot of example genes (rows) from Q1 with the highest tDE score in L2/3. Columns correspond to subclasses at P12 and P22, violins represent the expression distribution, and color represents median expression. Genes are colored according to the functional categories as in panel C. (**F**) FISH for tDE genes. Representative images (*top*) from an ‘across-row’ (see Methods) section in wS1. Cortical layers 2/3 and 4 are indicated on the sections. Quantification (*bottom*) of mean intensity in nuclei revealed strong temporal regulation of gene expression in L2/3 consistent with what was measured with snRNA-seq (see **Fig 2E**). Mann-Whitney test, ****p< 0.0001. L2/3_*Col19a1*: n = 4133, 3268 nuclei, L4_*Col19a1*: n = 1904, 1665 nuclei, L2/3_*Sorcs3*: n = 4414, 3394, L4_*Sorcs3*: n = 2221, 1876, L2/3_*Gabrg3*: n = 4340, 3210, L4_*Gabrg3*: n = 1808, 1564. Three mice per condition, two slices per mouse.

To understand the functional significance of Q1-Q4 for glutamatergic neurons, we computed the enrichment of curated lists of genes such as transcription factors (TFs), cell adhesion molecules (CAMs), ion channels (ICs), and housekeeping genes (HK) in glutamatergic subclasses using a Fisher’s exact test (**Fig 2C, Methods**). Housekeeping genes generally had low scores on both scales and were therefore enriched in Q3, consistent with their constitutive and non-specific expression. TFs were not significantly enriched in any quadrant, which is in line with their broad roles in regulating transcription throughout development across all subclasses (21,22). We found that CAMs, many of which are involved in regulating circuit formation (23), were enriched strongly in Q1 and, to a lesser degree, in Q4. This is consistent with their previously reported cell type-specific and dynamic expression patterns during circuit formation in other organisms, suggesting that this is a conserved feature of neurons during key phases of wiring (22,24–26). ICs were also enriched in Q1 and Q4, which aligns with changes in the physiological properties of neurons during this developmental period. Consistent with these trends, a Gene ontology (GO) analysis focusing on “biological process” revealed enriched categories such as “cell adhesion” and “axon guidance” for Q1 genes. In contrast, Q3 was enriched for general cellular processes such as “protein/mRNA transport” and “chromatin reorganization” (**Fig 2D, Fig S1D**). Q2 was enriched in “synaptic translation” terms, reflecting a global developmental effort to synthesize proteins, and it also had ∼60% of its enriched GO terms (8/13) overlapping with those of Q3 (**Fig S1D, S2A**). Q4 had a few terms overlapping with Q1 and many overlapping with Q3 (**Fig S2A**). Among its top terms were “ion transport”, “cell-cell adhesion”, and “protein phosphorylation” (**Fig S1D**).

Together, these results suggest that Q1 is a subclass-specific program related to circuit development, Q2 is a global developmental program, Q3 is a static “housekeeping” program, and Q4 is a subclass-specific program associated with neuronal identity. These trends were qualitatively similar in GABAergic subclasses except with fewer Q1 and Q2 genes (**Figs 2B, S2B**).

Next, we examined the degree to which temporally regulated genes were shared among subclasses. We isolated each subclass and identified significantly up/down-regulated genes (fold-change > 2 between P12 and P22, FDR < 0.05 Wilcoxon rank-sum test) (**Table S2**). Of the ∼421 genes identified, >60% were up/downregulated in a subclass-specific fashion (**Figs S2C-D**). These temporally regulated genes are a subset of the Q1-Q2 genes from **Fig 2B**. The subclass-specific genes were enriched in Q1, and the shared genes were restricted to Q2 **(Figs S2E,F).** Additionally, we detected ∼1.5x more downregulated than upregulated genes (**Fig S2D**). A similar pattern has been observed in multiple studies of fly brain development, where repression of gene expression is a dominant developmental driving force (27–29).

**Fig 2E** shows some examples of temporally regulated genes enriched in L2/3 glutamatergic neurons, including CAMs, TFs, ICs, and other genes. To confirm the temporal and subclass-specific expression patterns seen with snRNA-seq, we performed multiplexed fluorescence *in situ* hybridization (FISH) using the RNAscope assay (**Fig 2F**). First, we verified the RNAscope assay was working as intended using proprietary positive and negative control probes. As expected, the positive control probes showed specific and widespread expression in the brain, and the negative control probes revealed no detectable signal (**Fig S3E-F)**. Next, we selected three temporally regulated genes enriched in L2/3 from **Fig 2E** (*Col19a1*, *Sorcs3*, and *Gabrg3*), and probed their expression at P12 and P22 in wS1 (**Fig 2F**). The pattern of temporal regulation of gene expression was consistent between snRNA-seq and FISH (**Fig 2E-F**), validating our findings of subclass-specific temporal regulation of gene expression in S1 during postnatal development.

### Developmental changes in L2/3 cell type composition

We next analyzed the maturation of cell types between P12 and P22. Using a decision tree-based classifier (30) trained on the cell type transcriptional profiles at P22, we assigned P22 type labels to each P12 transcriptome (**Figs 3A, S3A-B; Methods**, **Table S3**). Based on transcriptomic similarity, each neuron at P12 could be unequivocally classified into one of the 45 cell types at P22. This mapping allowed us to compare each cell type between the two ages.

**Figure 3.**
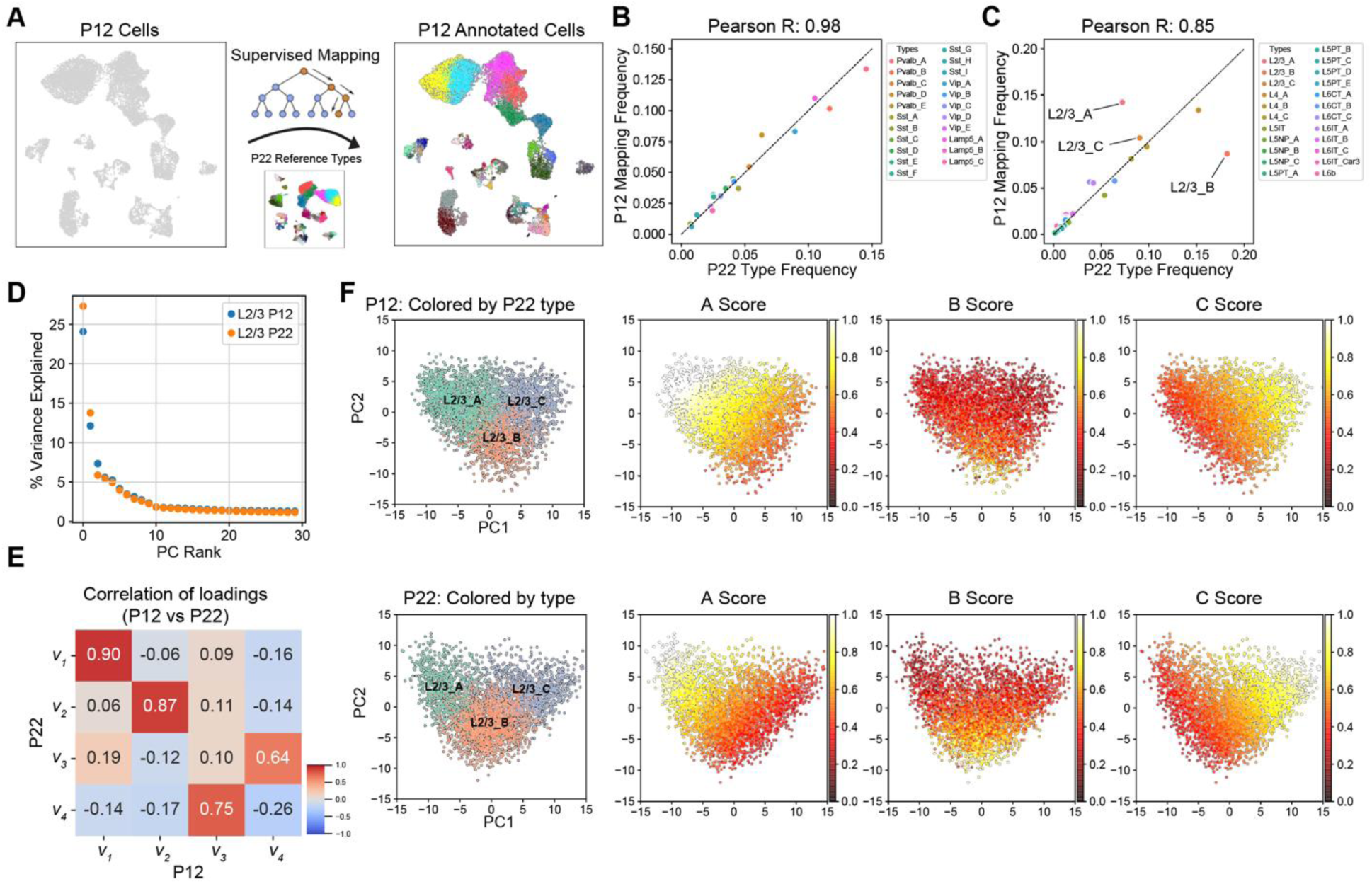
L2/3 pyramidal cell type composition is selectively regulated during development. (**A**) Schematic for transferring P22 cell type labels (reference data) to P12 cells to facilitate cell type-level comparisons. (**B**) Within GABAergic neurons (∼20% of all neurons), all cell types have approximately the same relative frequency between P12 (y-axis) and P22 (x-axis). Pearson correlation coefficient between the relative frequencies is indicated on top. (**C**) Same as panel B, for glutamatergic neurons (∼80% of all neurons). All glutamatergic cell types except for L2/3_A and L2/3_B have approximately the same relative frequency between P12 and P22. (**D**) Percent variance captured (y-axis) by each principal component (PC) within L2/3 neurons at P12 and P22 (colors). Note that PCA is performed independently on each dataset. For both ages, a spectral gap is observed after PC1 and PC2. (**E**) Pair-wise Pearson R values between the first four principal eigenvectors between P12 and P22. The first two principal eigenvectors corresponding to PC1 and PC2, which dominate the variance, map 1:1 between both ages with a high correlation. (**F**) Scatter plot of PC1 vs PC2 for L2/3 neurons at P12 (top row) and P22 (bottom row). With each row, the leftmost panel highlights cells by their type-identity (L2/3_A, L2/3_B, and L2/3_C). In the remaining panels within each row, cells are colored based on their aggregate expression levels of markers for each type (see **Methods**). Between P12 and P22, L2/3_A markers decrease in expression and increase in specificity, while L2/3_B markers increase in expression during development, driving cell type identity maturation.

Despite spanning two orders of magnitude, cell type frequencies tightly corresponded between P12 and P22 within all four GABAergic subclasses and 7/8 glutamatergic subclasses (**Fig 3B-C**). This suggests that most wS1 cell types acquire a distinct and stable transcriptional signature prior to P12. The only exception was L2/3 glutamatergic neurons (**Fig 3C**). Similar to what was observed in V1 (10), L2/3 glutamatergic neurons in wS1 can be clustered into three nominal types that we label L2/3_A, L2/3_B, and L2/3_C. The relative frequency of L2/3_C was similar between the two ages, but L2/3_A halved from P12 to P22, while L2/3_B doubled from P12 to P22 (**Fig 3C)**. Despite this prominent change in cell type composition, a principal component analysis (PCA) revealed that the transcriptomic programs underlying L2/3 heterogeneity were similar. The first two principal eigenvectors from PCA corresponded highly between P12 and P22 (**Fig 3D-E**). The correspondence was nearly 1:1 for the top two eigenvectors, but with a slight reduction in the correlation coefficient, likely due to the compositional shift of L2/3 neurons from P12 to P22. This temporal variation in cell type frequency coincided with i) a marked increase in the specificity of type A and C markers and ii) increased expression strength of B markers from P12 to P22 (**Figs 3F, S3C**). Consistent with the shift in cell type composition, >20% of L2/3 type-specific markers were temporally regulated, the highest among all subclasses (**Fig S3D**).

### Genes distinguishing L2/3 cell types and their temporal regulation

We sought to validate the cell type compositional shift described above by performing fluorescence *in situ* hybridization (FISH) in wS1. We targeted markers enriched in the three L2/3 types - *Cdh13* for L2/3_A, *Trpc6* for L2/3_B, and *Chrm2* for L2/3_C (**Fig 4A-B**). These three genes were reported earlier to mark L2/3 types in V1 (10). FISH experiments showed that between P12 and P22, *Cdh13+* L2/3_A cells decreased in frequency, *Trpc6+* L2/3_B cells increased in frequency, and *Chrm2+* L2/3_C were stable (**Fig 4C-D**). At P12, L2/3_A cells were concentrated at the L1/2 border, with a small population of L2/3_B cells and a large population of L2/3_C cells. At P22, there were very few L2/3_A cells at the L1/2 border, and the majority were concentrated in the middle of L2/3 (**Fig 4E**). Similar to what was observed in V1, we showed that *Cdh13+* cells in the center of L2/3 are likely inhibitory neurons, as they do not co-express *Slc17a7* (vGlut1) (**Fig S5A)**. Between P12 to P22, the *Chrm2+* L2/3_C cells maintain their frequency, whereas the *Trpc6+* L2/3_B cells increase relative to *Cdh13+* L2/3_A cells **(Fig 4E)**. In contrast to V1, where L2/3_A cells retain their L1/2 localization at ∼P21 despite decreasing in frequency, in wS1, L2/3_A cells are reduced at the L1/2 border while *Trpc6+* L2/3_B cells become more prevalent (**Fig 4E, Fig S5B-C**). As another validation of L2/3 compositional changes, we targeted *Adamts2* for L2/3_A, *Bdnf* for L2/3_B, and *Chrm2* for L2/3C (10,31). The expression of *Adamts2* and *Bdnf* showed the expected trend between P12 and P22 (**Fig S5D-E**), with *Adamts2+* cells decreasing in abundance while *Bdnf+* increased, agreeing with the snRNA-seq results (**Fig 4A**).

**Figure 4.**
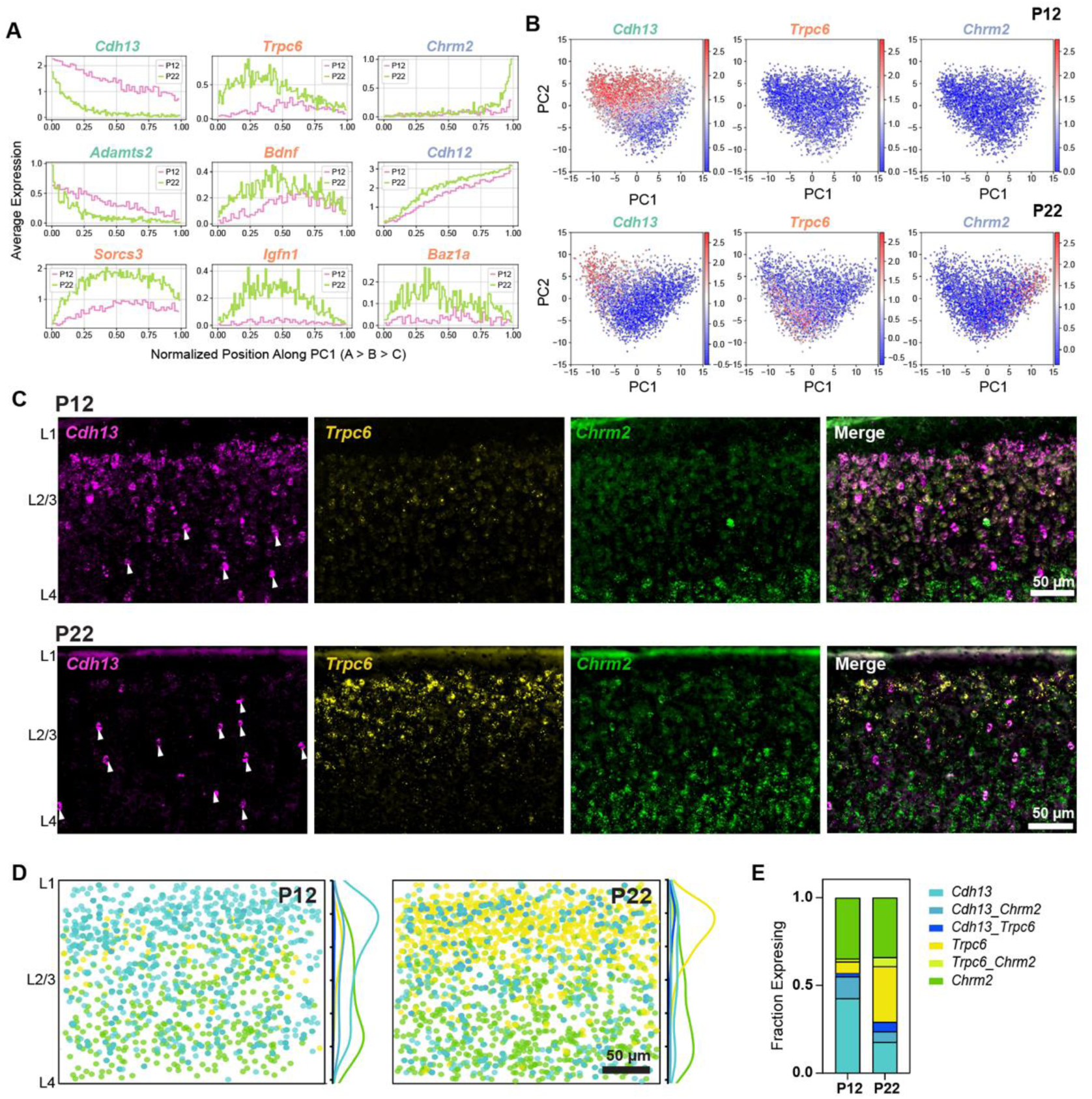
Cell type-specific temporal gene expression changes in L2/3 glutamatergic neurons. (**A**) Expression patterns of some L2/3 cell type-enriched genes along PC1 from **Fig 3F**. Genes are colored based on their type enrichment: A, green; B, orange; C, purple. Other genes are shown in **Fig S4**. (**B**) Same as **Fig 3F**, with cells colored by expression levels of *Cdh13* (left), *Trpc6* (middle), and *Chrm2* (right), which are targeted for FISH experiments in panels C-E. (**C**) Representative FISH images showing labeling of cell type markers *Cdh13, Trpc6,* and *Chrm2* at P12 and P22 within wS1 L2/3. Arrowheads indicate putative *Cdh13*-expressing interneurons (See **Fig S5A)**. (**D**) Summary plots based on overlay of all images of L2/3 visualizing expression of *Cdh13, Trpc6,* and *Chrm2* at P12 and P22. Circles represent individual excitatory cells within L2/3, colored based on their expression of one or more marker genes (color code as in panel E). To the right of each summary plot is the kernel density estimate for each type along the pial-ventricular axis. *Cdh13+* cells dominate in upper L2/3 at P12, whereas *Trpc6+* cells are more abundant at P22. N = 10-12 slices from 3 mice per time point. (**E**) Quantification of the fraction of excitatory (*Slc17a7+,* not shown) L2/3 cells expressing one or more markers *Cdh13*, *Trpc6*, and *Chrm2* at P12 and P22. N = 10-12 slices from 3 mice per time point.

Additionally, **Fig S4** highlights the temporal regulation of several TFs, ICs, and CAMs differentially expressed among L2/3_A-C types. Note that genes enriched within a given type do not perfectly overlap and often bleed into the adjacent type, and this is related to the continuous nature of transcriptional variation within L2/3 (10,11,32). Some genes that retain expression patterns from P12 to P22 (e.g., *Meis2*, *Foxp1, Kcnh7, Dscaml1*) may be involved in the initial establishment and/or maintenance of cell type identity. Other genes (e.g., *Rfx3*, *Zbtb16*, *Scn9a*, *Ncam2*) are significantly temporally regulated and may be involved in the refinement of these cell type identities, including the maturation of their circuitry and physiology.

Taken together, our results suggest that for most neuronal cell types in wS1, transcriptomic distinctions and relative composition are established prior to P12 and persist through the period of early sensory experience. Even though there are significant gene expression changes related to growth and maturation (**Fig 2**), these changes do not impact cell type identity. The exceptions to this rule were L2/3 types whose composition and type-specific gene signatures are significantly altered between P12 and P22. This mirrors the scenario reported earlier in V1 (10), raising the question of whether the two cortices also share global and subclass-specific gene expression programs related to maturation, which we now address.

### Differences and similarities between wS1 and V1 in cell type development and gene expression changes

The results thus far have highlighted multiple similarities between wS1 and V1 maturation. For example, both regions share the same transcriptomic subclasses and markers, L2/3 is selectively regulated by development in both, and both contain equivalent L2/3 cell types and markers (10). These similarities motivated a systematic comparison of developmental gene expression changes between wS1 and V1.

We identified tDE genes in V1 between P14 and P21, time points in the published data closest to those used in this study (**Table S4**). As in wS1, temporal gene regulation in V1 was predominantly subclass-specific (**Figs S6A-B**). Hypergeometric tests revealed that subclass-specific tDE genes were shared between wS1 and V1 for most subclasses (**Fig 5A**). For a few subclasses (e.g. inhibitory Sst, Vip and Lamp5 and excitatory L4 and L6b), there were too few tDE genes to conduct the analysis (**Figs S2C, S6A**). The shared subset of temporally regulated genes was enriched in GO terms related to neuronal development, including “GABAergic synapse”, “glutamatergic synapse”, “axon”, “nervous system development”, “axon guidance”, and other related programs (**Figs 5B, S6C**). Within L2/3, the overlapping genes were drawn from various functional categories, including CAMs, TFs, ICs, and NTRs (**Fig 5C**).

**Figure 5.**
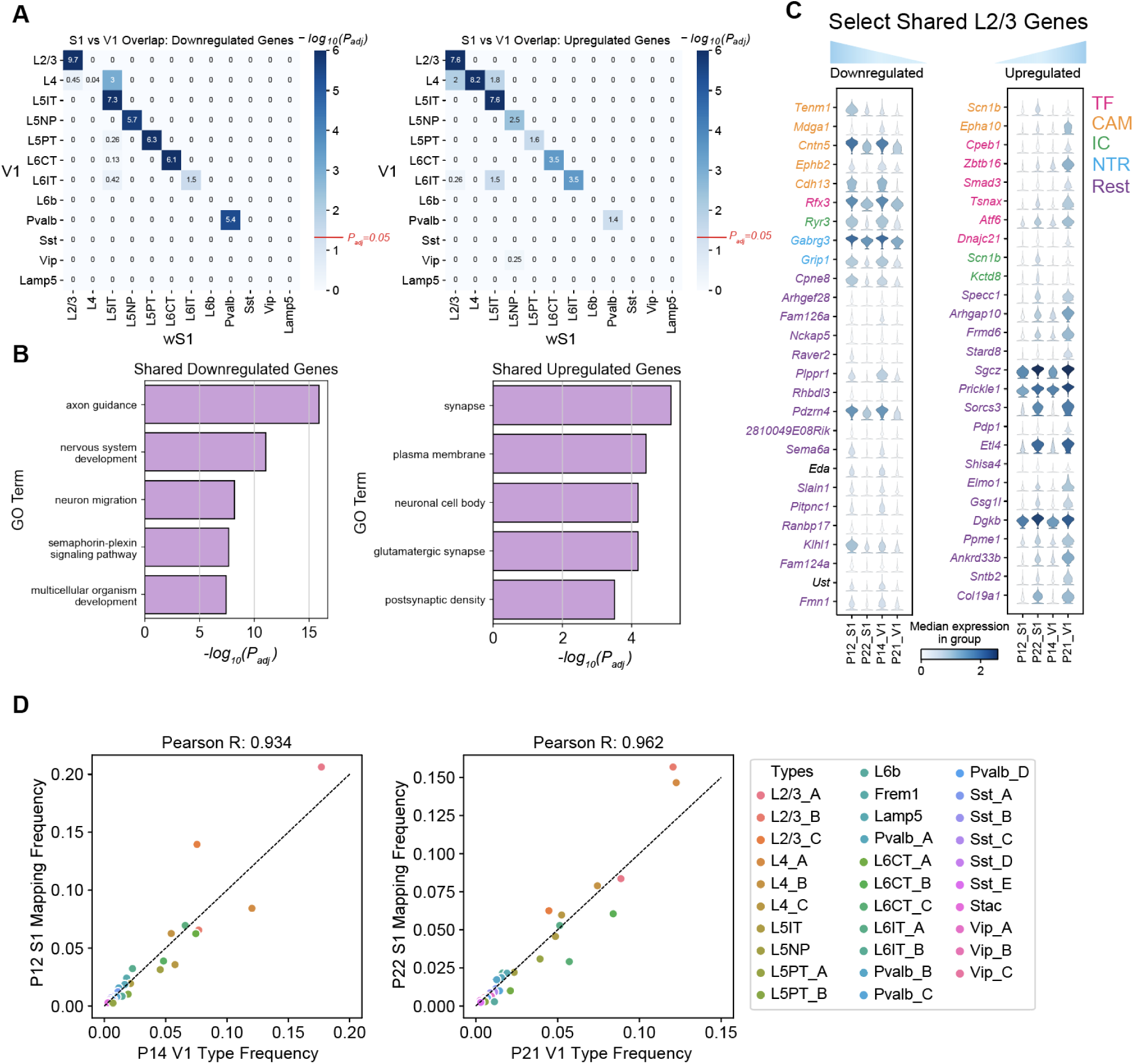
Comparative transcriptomic analysis of wS1 and V1. (**A**) Heatmap highlighting the overlap of tDE genes between all pairs of wS1 subclasses (P12 vs. P22; rows) and V1 subclasses (P14 vs. P21; columns). The left and right panels correspond to downregulated and upregulated genes. Values correspond to Bonferroni-corrected -log_10_(*P_adj_)* values from a hypergeometric test of overlap of tDE genes. The background set for these tests was the set of all the genes regulated in any subclass. Except for L4, subclasses with fewer than 10 tDE genes (**Figs S2C, S6A**) showed little to no overlap. The value *P_adj_* = 0.05 is highlighted on the scalebar (right). (**B**) Top 5 GO terms from shared downregulated and upregulated genes in V1 and wS1 in corresponding subclasses from panel A. See **Fig S6C** for a full list. (**C**) Examples of shared genes that are temporally downregulated (*left*) and upregulated (*right*) in L2/3 neurons between V1 and wS1. (**D**) Scatter plots showing highly similar relative frequencies between matched cell types (see **Fig S6D)** across V1 and wS1.

Moreover, wS1 and V1 share a striking correspondence in cell type composition. We trained classifiers on the V1 cell types at P14 and P21 and used them to transfer labels onto wS1 at P12 and P22, respectively. All 45 wS1 cell types mapped to the correct V1 subclass, and most cell types mapped 1:1 (**Fig S6D**). We also found that the relative cell type frequencies were highly concordant between the matched ages, indicating a high degree of overlap in cell type identity between the two cortical regions (**Fig 5D**). Together, this suggests that the two cortical regions share developmental programs at cell type resolution.

### Full-face whisker deprivation by plucking does not alter cell type development

In mouse V1, 1-2 weeks of dark rearing during development selectively alters the transcriptomes of L2/3 cell types, revealing a requirement of visual experience for cell type development (10,11). To test whether whisker deprivation has a similar effect on cell type development in wS1, we plucked all whiskers on the face from P12 to P22 and analyzed the resulting gene expression changes. We refer to this manipulation as 10-day all-whisker deprivation (10d AWD).

We repeated the analysis outlined in **Figs 3A-C**, training a classifier on the P22 cell types and using it to transfer P22 labels to 10d AWD transcriptomes. L2/3 composition and cell type frequency at P22 were minimally affected by 10d AWD (**Figs 6A-B**). Similar to the developmental analysis in **Fig 3**, we used PCA to study the influence on cell type composition in 10d AWD. For normal vs. 10d AWD P22, the top two principal eigenvectors, which captured >40% of the variance, exhibited a more robust 1:1 mapping compared to the case of the normal P22 vs. normal P12 (**Figs 6C,D, 3E**), consistent with 10d AWD having a minimal effect on the overall expression levels and the correlation structure of gene expression. Furthermore, the expression patterns of L2/3 type markers along the first two principal components (PCs) were highly similar between normal and 10d AWD mice at P22 (**Figs 6E-F, S7A**).

**Figure 6.**
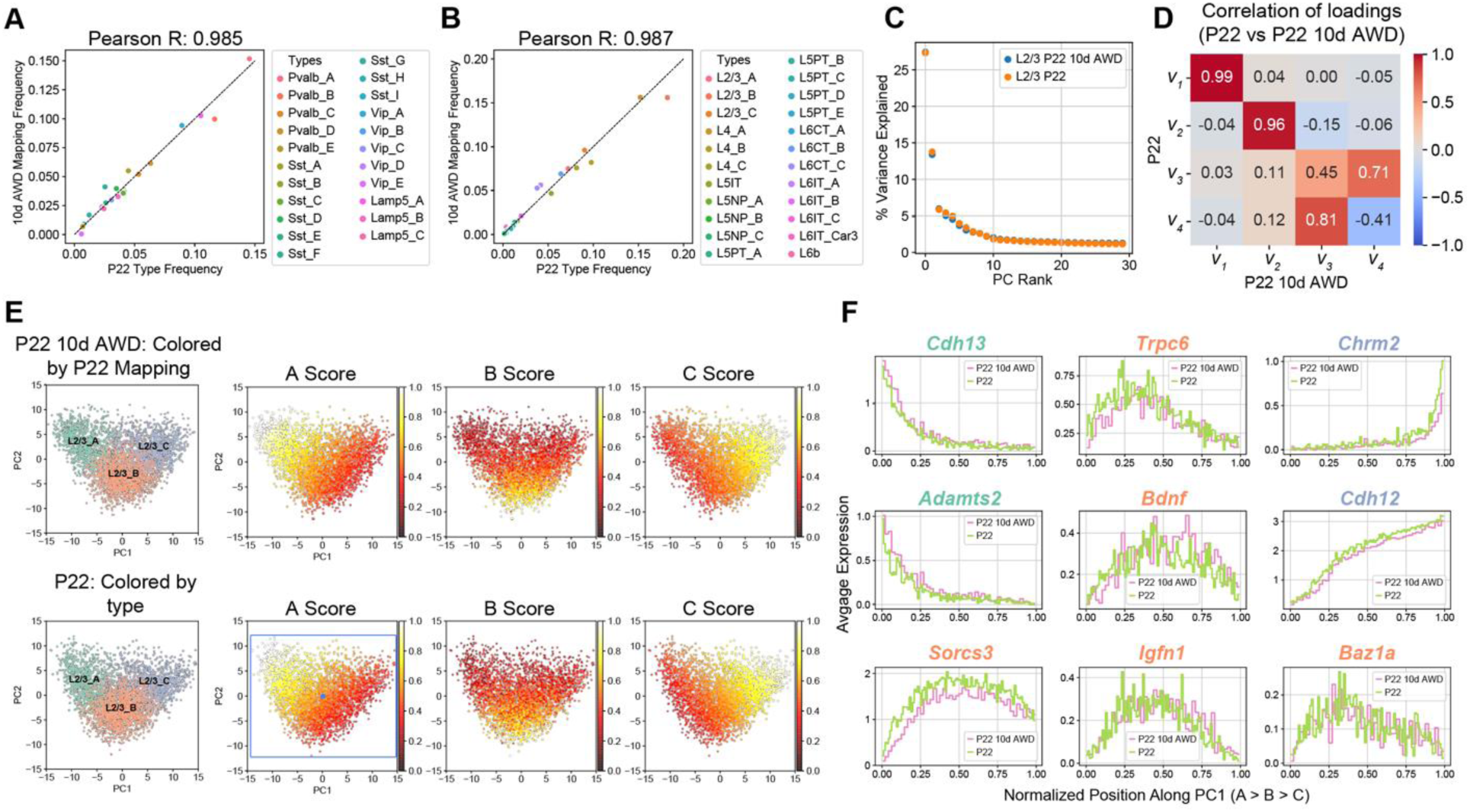
Full-face 10-day all-whisker deprivation (10d AWD) does not influence cell type maturation. (**A**) GABAergic cell types have approximately the same relative frequency in P22 10d AWD (y-axis) and normal P22 (x-axis). Note that cell type frequencies are normalized within all GABAergic neurons (∼20% of all neurons). (**B**) Same as panel A for glutamatergic neurons (∼80% of all neurons), highlighting similar cell type frequencies between P22 10d AWD (y-axis) and normal P22 (x-axis). (**C**) Similar to **Fig 3D**, showing that PC1 and PC2 are sufficient to describe transcriptional variance within L2/3 in the normal P22 and P22 10d AWD datasets. (**D**) Heatmap of Pearson correlation between the principal eigenvectors (as in **Fig 3E**) showing that the first two principal eigenvectors map 1:1 between the two datasets. (**E**) PC1 vs. PC2 scatter plot for L2/3 neurons at P22 10d AWD (top) and normal P22 (bottom), highlighting the location of types and the type-specific marker scores. Representation as in **Fig 3F**. Scores are similar between the two datasets (see also **Fig S7A**). (**F**) L2/3 markers genes, as in **Fig 4A**, are shown as a function of cells’ position along PC1 comparing patterns between normal P22 and P22 10d AWD.

Analysis of differentially expressed genes (DEGs) further confirmed that 10d AWD had a minimal influence on gene expression in all subclasses (**Table S5**). We found 51 DEGs, which is ∼8x smaller than the number of temporally regulated DEGs between P12 and P22 (**Figs S7B-D**). For most of DEGs, regulation was subclass-specific, with *Lamp5+* interneurons exhibiting the largest number of changes. Altogether, we conclude that full-face whisker deprivation between P12 and P22 minimally impacts the maturation of cell types and subclass-specific transcriptional programs in wS1.

### Brief row-based whisker deprivation upregulates column-specific activity-dependent gene expression programs in wS1

Previous work has shown that depriving a single row of whiskers for one day induces rapid plasticity in excitatory and inhibitory circuits in the columns corresponding to the deprived whiskers. This includes Hebbian and homeostatic synaptic plasticity and changes in the intrinsic excitability of neurons, with different mechanisms at play depending on the cell type and cortical layer (18,19,33–35). However, the molecular factors behind these layer-specific changes in response to whisker deprivation in are still unknown.

We performed 1-day deprivation of the B and D whisker rows (1d RWD), a manipulation that induces competitive whisker map plasticity (18), and assessed transcriptomic cell types and gene expression changes in S1 using snRNA-seq and FISH (**Fig 7A**). Cell type distributions were highly correlated between normal and 1d RWD datasets at P22 (**Figs S8A-B**). As in the normal and 10d AWD P22 datasets, variation within L2/3 neurons was captured well by the top two PCs (**Fig S8C**), which correlated 1:1 with the top two PCs of normal P22 data (**Fig S8D**). Furthermore, the expression patterns of L2/3 type markers along the first two principal components (PCs) were highly similar between normal and 1d RWD mice at P22 (**Fig S8E-G**). Altogether, this suggests that 1d RWD has a minimal influence on cell type identity in wS1, as expected for such a brief sensory manipulation.

**Figure 7.**
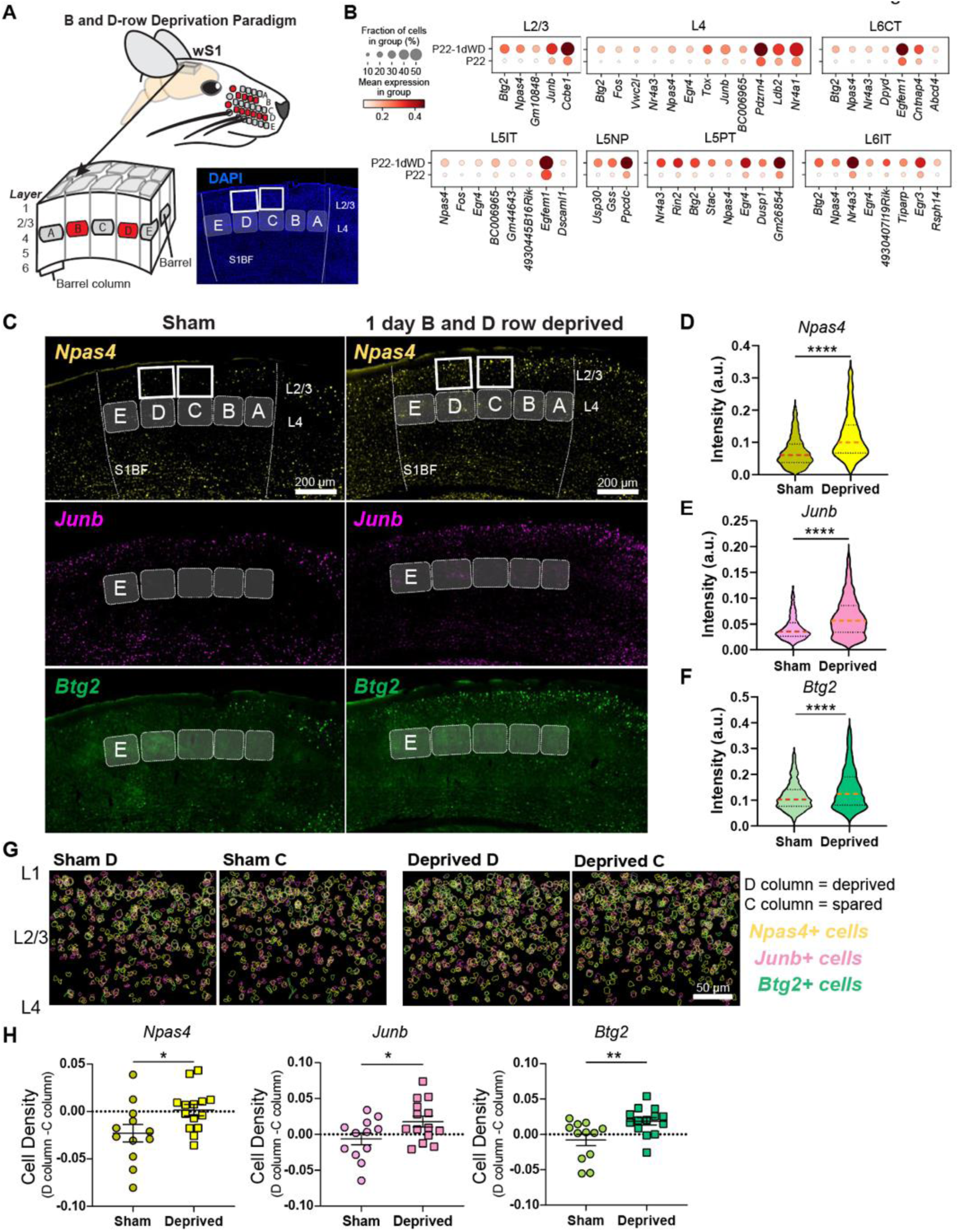
Brief row whisker deprivation (1d RWD) upregulates activity-dependent gene expression programs in deprived columns. (**A**) Schematic representation of the 1d row-whisker deprivation (RWD) manipulation and ‘across-row’ S1 section in which all barrel columns are identifiable. A representative image of DAPI labeling in ‘across-row’ S1 section. (**B**) Dotplots of genes upregulated by 1d RWD in glutamatergic subclasses (panels). Within each panel, rows indicate condition and columns indicate genes. The size of each circle corresponds to the % of cells with nonzero expression, and the color indicates average expression level. (**C**) Representative widefield images of wS1. Barrels and barrel fields are indicated with light grey rectangles and labeled. Boxes indicate the locations of ROIs used for analysis in **G.** (**D**) *Npas4* intensity inside *Slc17a7*+ L2/3 excitatory cells is increased after 1d RWD. Violin plots show median (dashed line) and quartiles (dotted lines). Mann-Whitney test, p < 0.0001 ****, n = 1571, 1337 cells respectively, from 3-4 mice per group. (**E**) *Junb* intensity inside *Slc17a7*+ L2/3 cells is increased after 1d RWD. Mann-Whitney test, p < 0.0001 ****, n = 1483, 1405 cells respectively, from 3-4 mice per group. (**F**) *Btg2* intensity inside *Slc17a7*+ L2/3 cells is increased after 1d RWD. Mann-Whitney test, p < 0.0001 ****, n = 1561, 1369 cells respectively, from 3-4 mice per group. (**G**) L2/3 C and D columns were compared to examine whether effects are specific to the deprived (D) column. Pseudocolored outlines of *Npas4, Junb, and Btg2* expressing cells. Each plot is an overlay of 5 images of L2/3 S1 from 3 mice. (**H**) Quantification of column-specific changes in 1d RWD upregulated genes measuring. Plotted values are the difference between the fraction of *Npas4/Junb/Btg2* expressing cells among all excitatory (*Slc17a7*+) cells in the L2/3 D column and the corresponding fraction in the L2/3 C column. Symbols represent slices from sham and 1d RWD mice. For each condition, mean and SEM (error bars) are shown. *Npas4*: Unpaired t-test, *p = 0.0321, n = 12,14 slices, 3-4 mice. *JunB*: Unpaired t-test, *p = 0.0427, n = 12,14 slices, 3-4 mice. *Btg2*: Mann-Whitney, **p = 0.0077, n = 12,14 slices, 3-4 mice.

It is well-established that sensory manipulation induces activity-dependent gene expression programs that result in long-lasting cellular adaptations essential for learning and maintaining circuit homeostasis (36,37). We hypothesized that 1d RWD may elicit an activity-dependent transcriptional response (**Fig 7A**). Within each neuronal subclass, we sought genes expressed in at least 20% of cells that were upregulated in 1d RWD mice (fold-change>2, FDR<0.05, Wilcoxon rank-sum test) (**Table S6**). We detected 59 such genes across all 12 subclasses. **Fig 7B** shows that the upregulated genes contain well-known “activity-dependent” genes such as *Npas4*, *Btg2*, *Junb*, and *Nr4a1* (36). Importantly, these genes were different from the few regulated by 10d WD (**Fig S7**), indicating their specificity to 1d RWD plasticity.

We performed FISH experiments to confirm these gene expression changes and assess their laminar and columnar organization in the barrel field (**Fig 7C**). We used a plane of section that contains one whisker column in each of the five rows (A-E), enabling whisker row identity to be recognized since each whisker row maps to an array of cortical columns in the barrel field (38) (**Fig 7A**). We probed for three IEG candidates shown in **Fig 7C**--*Npas4*, *Btg2*, and *Junb*. These mRNAs were significantly increased in L2/3 of wS1 as measured by intensity per L2/3 cell (**Fig 7D-F**). To determine if the mRNAs were upregulated in a whisker column-specific manner, we separately analyzed the number of cells expressing each mRNA in excitatory L2/3 neurons in the deprived D vs. non-deprived C column of the same tissue section from sham and deprived mice (**Figs 7G,H**). We found that all mRNAs were upregulated in D compared to C in the whisker-deprived but not in sham controls, highlighting whisker column-specific gene regulation during competitive map plasticity (**Fig 7H**). Together, these results suggest that 1d RWD triggers rapid and selective induction of genes functionally associated with activity-dependent plasticity in a spatially localized pattern wherein mRNAs are more strongly upregulated in the deprived columns than in neighboring spared columns. Such gene expression patterns may constitute an early molecular mechanism of competitive map plasticity.

## DISCUSSION

### A postnatal developmental atlas of mouse wS1

The somatosensory cortex is a canonical model to study experience-dependent and -independent development. Indeed, the principles of somatosensory organization and plasticity are similar between rodents and humans. Understanding the molecular programs underlying postnatal development of sensory cortex may be critical to understanding neurodevelopmental disorders such as autism that are associated with circuit dysfunction and impaired sensory processing. Prior studies have profiled the transcriptomic diversity of cell types in wS1 using scRNA-seq in adult mice (7) and bulk RNA-seq in adolescent mice (39–41), but the gene expression changes that guide the emergence of cell type identity during early postnatal development in wS1 have not been explored.

We used snRNA-seq to study the postnatal development of transcriptomic cell types in wS1 from P12 to P22, the period of juvenile development immediately following the onset of whisking. Our mouse wS1 atlas comprises ∼80,000 single-nucleus transcriptomes at two postnatal ages and across three rearing conditions. The overall taxonomy and frequency distribution of ∼45 neuronal cell types were highly conserved across the time points and rearing conditions, allowing us to disambiguate broadly shared from cell type-specific changes during normal development and in response to whisker deprivation.

Proper assembly of cortical circuits involves precise control of gene expression in space and time. The comparison of expression profiles between P12 and P22 revealed hundreds of temporally regulated genes in wS1, over 60% of which were subclass-specific and enriched for functionally relevant processes associated with neuronal maturation, such as axon guidance and synapse maturation. Overall, glutamatergic neurons contained a higher proportion of temporally regulated genes than GABAergic neurons. A recent comparative transcriptomics study of V1 across human, macaque, and mouse showed that excitatory neurons are more evolutionarily divergent than inhibitory neurons (42), similar to the increased excitatory neuron developmental regulation we observed in wS1. The upstream mechanisms controlling the timing and subclass-specific expression of these genes and their potential roles in circuit formation and refinement are important areas for future research.

### L2/3 cell types undergo significant changes during normal development

Of the 45 molecularly defined neuronal cell types, 42 were present at identical frequencies at P12 and P22, indicating that most wS1 neuron types are defined before P12. The only cell types that undergo significant changes in composition and cell type-specific gene expression between P12 and P22 are pyramidal cell subtypes in L2/3 (L2/3_A-C). The molecular identity and developmental changes of the three L2/3 cell types in wS1 mirror those observed in V1 (10), where L2/3 cell types have been shown to possess unique binocular tuning properties and correlate with distinct projections to higher visual areas (HVAs)(10,43). It is, therefore, reasonable to surmise that the three L2/3 cell types in wS1 may possess distinct functional properties and connectivity patterns. For example, *Cdh13* and *Cdh12* were expressed in different L2/3 neurons, with *Cdh13* enriched in type A and *Cdh12* in types B/C. A recent study demonstrated that *Cdh12+* and *Cdh13+* L5 pyramidal neurons in S1 are inhibited by CCK+ and PV+ interneurons, respectively (44). Based on these trends, it is likely that L2/3_A cells are inhibited by PV+ neurons and L2/3_B/C cells are inhibited by CCK+ neurons. Another study identified that a group of mouse S1 L2/3 cells enriched in the *Baz1a* gene (likely the L2/3_B type) adapt to changes in whisker input via transient changes in IEG expression (45). Thus, the delayed maturation of these L2/3 cell types may be linked to their experience-dependent plasticity in adulthood. Since the L2/3 cell types differ in laminar position, they may receive different thalamic inputs, with L2-situated cells (L2/3_A and B here) tending to receive less VPM axon innervation than L3-situated cells (L2/3_C)(46,47), as well as different local inhibitory inputs (48).

A potential mechanism underlying the developmental flexibility of L2/3 neurons could be the presence of a transcriptomic gradient. In principal component space, we observed that L2/3 neurons form a triangle, with each of the three cell types occupying a vertex. Additionally, many markers were expressed in a gradient fashion, like the functional gradients observed for L2/3 neurons in V1 (49). This suggests that wS1 L2/3 neurons exhibit an overall gradient pattern while also containing some specialist cell types, mirroring what has been observed in V1 and the intestine (10,11,50). Occupying a flexible continuum could facilitate the drastic gene expression and compositional shifts observed in these neurons from P12 to P22.

### Developmental transcriptomic conservation and divergence between V1 and wS1

Recent studies have shown a high degree of correspondence in neural diversity and cell type markers between cortical areas (51,52), and this was indeed the case between wS1 and V1. Our comparison of P12 to P22 in wS1 and P14 to P21 in V1 revealed similar developmental gene expression patterns. In most cases, there was significant overlap in the up- and down-regulated genes between matched neuronal subclasses between the two regions and little to no overlap across distinct subclasses. The notable exception was that L4 neurons did not share downregulated genes between the two regions (**Fig 6A**), which may be related to the fact that L4 contains spiny stellate cells in wS1 but not V1.

Despite this broad similarity, there was a laminar difference in the organization of L2/3 neuron subclasses between the two regions. Our data suggests that wS1 undergoes a similar remodeling of cell types from P12 to P22 as V1, with A-like cells decreasing and B-like cells increasing in number. However, unlike V1 where the L2/3_A-C cells segregate into sublaminae along the radial (pial-ventricular) axis at P21, we found that, in wS1, L2/3_A and L2/3_B somas are intermixed in superficial L2/3, but are spatially separated from the C somas, which reside in deep L2/3. The factors driving this difference between V1 and wS1 are unknown but may stem from broader differences in laminae across the sensory cortices (53).

The overall similarity in gene expression programs appears reasonable, given that V1 and wS1 share many aspects in the timing of their postnatal development. Thalamocortical axons begin innervating the cortical plate and reaching their targets in L4 around the same time in wS1 and V1 (54,55). Callosal axons cross the midline around the same time in V1 and wS1 also and undergo activity-dependent rearrangement during postnatal week two. By the second postnatal week, sensory input increases in V1 and wS1 at the onset of activity whisking (P12) and eye-opening (P14). However, an important difference is that the maturation of wS1 is accelerated compared to V1. Sensory-evoked responses can be elicited even before birth with whisker stimulation, in contrast to V1 where these responses arise by the end of the first postnatal week. The classically defined critical periods for whisker deprivation-induced plasticity in wS1 close in postnatal weeks 1-2, while in V1 the classical critical period for ocular dominance plasticity doesn’t end until P32 (56).

Despite the robust effects of whisker experience on wS1 circuit function, our results demonstrate that normal whisker experience from P12 to P22 is not required for the development of cell types in wS1. This contrasts with our previous observations in V1, where dark rearing led to significant changes in gene expression and composition of L2/3 cell types, resulting in altered spatial organization and functional tuning (10,11). In wS1, 10d AWD did not impact gene expression or composition of L2/3 cell types. This result suggests that sensory experience may vary in importance for the maturation of cortical cell types across regions. In wS1, the proper formation and stabilization of neural circuits may rely more heavily on intrinsic programs, regardless of sensory input (57).

Why might L2/3 cell type development be regulated by visual experience in V1, but not by 10d AWD whisker deprivation in wS1? One possibility is that the overall impact of sensory experience on cortical cell and circuit development is milder in wS1 than in V1. Consistent with this idea, ocular dominance shifts of V1 neurons driven by monocular deprivation are often stronger than whisker tuning shifts in wS1 driven by plucking a subset of whiskers (58,59). During the critical period for ocular dominance plasticity in V1, visual experience shapes thalamocortical axons. It is required to maintain their structure, while in wS1, whisker sensory use does not alter thalamocortical or L4 topography after P4, and before that only nerve or follicle damage is sufficient to drive plasticity (55,60). The critical period for ocular dominance plasticity has a stark ending at P32 in mice (56), while S1 retains many forms of experience-dependent plasticity into adulthood (5), suggesting that wS1 plasticity may not be as tied to developmental mechanisms as in V1.

Alternatively, the lack of effect of whisker deprivation on wS1 cell type development may stem from differences in the efficiency of whisker vs. visual deprivation paradigms (13). Visual deprivation, such as dark rearing, completely removes sensory drive and statistical patterns of sensory input, leaving only retinal spontaneous activity, which has different properties. In contrast, whisker plucking does not fully eliminate external sensory input because direct contact with the skin of the mystacial pad, e.g. during grooming or cuddling, still provides some afferent activation. Thus, plucking is likely to eliminate less sensory input than dark rearing and, hence, may have a less pronounced impact on S1 gene expression.

### Competitive whisker deprivation drives plasticity-related gene expression programs in L2/3

Despite the lack of requirement for whisker experience on cell type maturation, we found that brief whisker deprivation of only two rows of whiskers (1d RWD) upregulates activity-dependent gene expression programs in multiple wS1 cell types, including *Npas4, Btg2, Junb, and Nr4a1*, which are known to be involved in synaptic plasticity and circuit homeostasis (36). Typically, expression of these IEGs is thought to be driven by increased levels of activity, as occurs during environmental enrichment (P65-75 animals) in S1 (61,62) or dark rearing followed by light exposure in V1 (63). A recent study in V1 showed that seven days of dark-rearing followed by 1- 2 hours of light exposure induced IEG expression (63). Interestingly, dark-rearing alone also increased the expression of several genes, particularly in L2/3 excitatory neurons (63). Our finding that a subset of IEGs are upregulated in response to deprivation indicates that IEG induction may represent a general plasticity program broadly used to engage plasticity mechanisms rather than a specific consequence of elevated neural activity. Notably, these IEGs were expressed mainly in the L2/3_B cell population, highlighting the role of L2/3_B cells as a critical functionally distinct subpopulation in L2/3 (45).

We found that the IEGs were upregulated in a column-specific fashion in wS1. Deprived columns showed strong upregulation of IEGs relative to neighboring spared columns. This deprivation paradigm triggers synaptic plasticity, a component of competitive map plasticity, wherein whisker-evoked responses to deprived whiskers weaken and shrink within the whisker map while responses to spared whiskers strengthen and expand (64). Thus, the selective upregulation of IEGs in deprived columns may represent an early molecular step underlying competitive map plasticity. Together, the 1d RWD results indicate that whisker experience exerts only relatively subtle effects on the development of cell types, but has significant effects at the gene expression level within specific cell types.

### Summary and limitations

This work contributes a single-nucleus transcriptomic resource for postnatal wS1 development under normal and deprived sensory experience. We delineated developmental changes in neuronal subclasses and types, and found that these were highly conserved between mouse V1 and wS1. However, whisker experience served a limited role in wS1 cell type development, which contrasts with the greater role of vision in V1 cell type development. Nevertheless, brief whisker deprivation induced significant changes in the expression of IEGs in a whisker column-specific fashion. Thus, while wS1 may rely less on experience and more on hardwired genetic programs than V1 to develop cell types, these cell types can undergo experience-dependent gene expression changes.

Throughout the study, we validated multiple scRNA-seq findings by performing FISH experiments targeting representative genes. Future studies could quantify the expression of many genes simultaneously in the same cell as a function of the cell’s location, especially in the context of wS1 barrels, using spatial transcriptomics approaches to further test the hypotheses established in this work. The gene regulatory logic that translates experience-driven activity to cell-type specific transcriptome changes remains poorly explored. Further research aimed at dissecting patterns at the DNA level, such as chromatin organization and DNA methylation, could reveal the gene regulatory networks responsible for the transcriptomic patterns observed here. We also note that the paucity of changes in the transcriptome during 10d AWD or 1d RWD may be due to low sampling resolution in snRNA-seq, or because the underlying changes could be at the protein level. Finally, additional work could relate the cell type-specific IEG expression during 1d RWD reported here to changes in neuronal physiology, for example, by labeling specific cell types and surveying their properties using slice electrophysiology or by performing 2P imaging of labeled cell types *in vivo* during whisker sensory behavior.

## MATERIALS AND METHODS

### Computational Methods

#### Alignment and gene expression quantification

FASTQ files with raw reads were processed using Cell Ranger v7.0.1 (10X Genomics) with default parameters. We used the mm10 (GENCODE vM23/Ensembl 98) reference genome and transcriptome. Reads aligning to the entire gene body (exons, introns, and UTRs) are used to quantify expression levels. Each single-nucleus library was processed using the same settings to yield a gene expression matrix (GEM) of mRNA counts across genes (columns) and nuclei (rows). Each row ID was tagged with the sample metadata for later meta-analysis. We henceforth refer to each single-nucleus transcriptome as a ‘‘cell.’’

#### Initial pre-processing and quality control of snRNA-seq data

This section outlines the initial transcriptomic analysis of data from all samples. Unless otherwise noted, all analyses were performed in Python using the SCANPY package(65) based on the following steps https://github.com/shekharlab/wS1dev:

1. Raw GEMs from 12 snRNA-seq libraries were combined: P12 control (A1, A2), P22 control (S1, S2, S3, S4), P22 1d RWD (D1, D2, D3, D4), and P22 10d AWD (C1,C2). This resulted in a GEM containing 199,524 cells and 32,285 genes.
2. We generated scatter plots of the number of transcript molecules in each cell (n_counts_), the percent of transcripts derived from mitochondrially encoded genes (percent_mito), and the number of expressed genes (n_genes_) to identify outlier cells. Cells that satisfied n_genes_ < 7000 and n_counts_ < 40,000, and n_counts_ > 500 were retained, and genes detected in fewer than 100 cells were removed from further analysis. This resulted in a GEM of 199,074 cells and 21,258 genes (**Fig S1A**). Cells were normalized for library size differences by rescaling the transcript counts in each cell to a total of 10,000, followed by log transformation.
3. For clustering and visualization, we followed steps described previously(10). Briefly, we identified highly variable genes (HVGs), z-scored expression values for each HVG across cells, and used the z-scored data to compute a reduced dimensional representation based on principal component analysis (PCA). The top 40 principal components (PCs) were used to build a nearest-neighbor graph on the cells. We clustered this graph using the Leiden algorithm(66) and embedded it in 2D via the Uniform Manifold Approximation and Projection (UMAP) algorithm(67). We identified 39 preliminary clusters.
4. Four of the 39 preliminary clusters contained 20-40% mitochondrial transcripts, while the remaining clusters contained, on average, 5-10% mitochondrial transcripts. We removed these clusters from the data, resulting in 169,674 cells (**Fig S1A**). One cluster belonged to samples S1/S2, another to A1/A2, another to C1/C2/S4, and the last one to S3/D3/D4.
5. With this filtered set of 169,674 cells, we re-ran the clustering pipeline described above to obtain a new set of clusters. To assess the quality of these clusters, we trained a multi-class classifier using XGBoost, a gradient boosted decision tree-based classification algorithm(30) on the subclasses from the primary visual cortex(10) (V1). From the V1 study, we used all of the postnatal time points collected from normally reared animals. 28 out of the 39 preliminary clusters mapped strongly to one of the 20 V1 subclasses. However, 11 clusters mapped diffusely to the subclasses and/or mapped with low classifier confidence. Further inspection indicated that the poorly mapping clusters had higher doublet scores than tightly mapping clusters(20), and their top markers were enriched in gene ontology (GO) terms related to “apoptosis” and “response to toxic substance”. Removal of the 11 problematic clusters reduced the number of cells to 114,812. Next, we examined the V1 subclass composition of each remaining cluster and removed cells corresponding to any V1 subclass that accounted for <2% of the cluster. This further purified clusters, ultimately yielding 112,233 cells (**Fig S1A**).
6. Finally, we isolated each of the four experimental conditions (P12 control, P22 control, P22 1d RWD, and P22 10d AWD) and re-ran the clustering pipeline described above. Within each condition, we removed clusters containing cells that mapped to V1 subclasses in a non-specific manner. This yielded a final number of 111,299 cells that form the foundation of this paper (**Fig S1A**). Only the 81,462 neurons were used in all the analyses reported in this paper.

#### Temporal regulation and subclass specificity analysis

The gene quadrant analysis (**Fig 2)** was performed on the P12 and P22 control datasets separately for the glutamatergic and GABAergic neuronal classes. Within either class, we performed a Wilcoxon rank-sum test for each subclass against the rest of the subclasses to identify subclass-specific markers (fold-change (FC)>2, FDR<0.05, expressed in >40% of cells). For each pair of gene and subclass, we calculated fold changes separately at P12 and P22 and selected the maximum value as the subclass variability (SV) score. We next performed Wilcoxon rank-sum test between P12 and P22 for each subclass to identify temporally regulated genes (FC>2, FDR<0.05, expressed in >40% of cells). The maximum temporal FC value for each gene across all subclasses was assigned to be its tDE score. To define the quadrants, a threshold FC value of 2 was chosen, and we verified that values between 1.5 and 2.5 do not qualitatively impact the results shown in **Figs 2** and **S2**.

In **Figs 2B,C** of the main text, we assess the statistical enrichment of specific gene groups in each quadrant using a Fisher’s Exact Test. For each quadrant, we calculated the odds ratio (OR) and p-value using Fisher’s Exact Test to determine whether TFs, CAMs, ICs, or HKs were significantly over- or under-represented compared to the null expectation based on all genes. Given the multiple comparisons, we applied the Bonferroni correction to adjust the p-values, controlling for false positive rate. The results were visualized by categorizing the quadrants based on statistically significant under- or over-representation: quadrants were shaded grey if the adjusted p-value exceeded 0.05 (no enrichment), red if the adjusted p-value was less than 0.05 with OR < 1 (significant under-representation), and green if the adjusted p-value was less than 0.05 with OR > 1 (significant over-representation). This approach allowed us to identify statistically significant enrichment patterns, highlighting their potential regulatory roles in wS1 development. Finally, gene ontology (GO) enrichment analysis was performed on each quadrant’s set of genes using the python package GOATOOLS(68). We used the default background set in GOATOOLS, which comprised all protein-coding genes.

#### Subclass-by-subclass differential gene expression analysis between each experimental condition and P22 control

We sought to identify gene expression differences at the subclass level between the P22 control dataset and the three experimental datasets, including P12, P22 10d AWD, and P22 1d RWD. Here, each subclass was isolated, and a Wilcoxon rank-sum test was performed for that subclass between P22 control and each of the three conditions. Of the genes expressed in >40% of cells in one of the two datasets, we selected those with FC>2 at false-discovery rate (FDR)<0.05. A python implementation of the R package UpSetPlot (69) was used to compute and visualize the number of genes regulated in a shared and subclass-specific fashion. For these differential gene expression tests, samples S3, D3, and D4 were excluded due to their relatively low number of transcript counts and expressed genes to not affect the results (**Fig S1B-C**). However, these samples were not excluded from any other analysis and consistently contained all neuronal subpopulations with no frequency discrepancies (**Fig 1F**).

#### Class, subclass, and cell type annotation of snRNA-seq data

We annotated cells in our dataset according to the taxonomy in **Fig 1D**. The two major neuronal classes were easily identified using glutamatergic marker *Slc17a7I* and GABAergic markers *Gad1* and *Gad2* in each condition separately. Additionally, we identified non-neuronal groups using known markers(31). Non-neuronal cells were discarded at an early stage, and we focused on the two neuronal classes, which clustered distantly from each other in gene expression space. Clusters within each condition naturally separated according to subclasses that could be annotated based on established markers(10). The class and subclass levels of the taxonomy are highly conserved across cortical regions and studies. However, as the cell type level tends to vary across regions, studies, and conditions, we annotated cell types in the P22 control dataset, and then transferred these labels onto the other conditions.

To annotate cell types in the P22 control dataset, we first trained a classifier on the cell types of the P21 V1 dataset(10). We then applied the classifier to the P22 control dataset to assign each wS1 neuron a P21 V1 cell type label. Second, we isolated each subclass and used Leiden clustering(66), varying the resolution parameter from 0.25 to 2, with higher values providing more clusters. Third, we trained and validated a classifier for each clustering resolution, identifying the resolution at which cluster validation error (computed using held-out cells) increases significantly, indicating over-clustering. The general procedure to train and validate such classifiers is described in the following section. Another telltale sign for diagnosing over-clustering was when differential expression analysis yielded highly overlapping marker sets for their clusters. Ultimately, these steps nominated a range of optimal clustering resolutions, and we chose the final resolution at which the validation error was low (90%) and there was a high concordance with the V1 clustering. This resolution also yielded unique marker sets across the cell types.

Finally, we combined all subclasses for each of the two classes (glutamatergic and GABAergic neurons), and used a classification analysis to verify that we have not over-clustered the data. We trained a classifier on the P22 wS1 control data and mapped the P21 V1 data to it. Clusters from the P22 wS1 data were removed if they satisfied the following criteria simultaneously: 1) received no mapping from V1, 2) had a high doublet and/or mitochondrial score, and 3) were not learnable during training. This procedure filtered ∼800 cells, approximately 2.5% of the P22 control data.

#### Classifier-based mapping of experimental conditions to P22 control data

To assess transcriptomic correspondence of clusters across ages (P12 vs. P22) or between rearing conditions (control vs. 1d RWD and 10d AWD), we used XGBoost, a gradient boosted decision tree-based classification algorithm(30). These analyses were performed to study the effects of development (P12), sensory experience (P22 10d WD), and rapid homeostatic plasticity (P22 1d WD) on cell type identity and composition. We also used this approach to compare the cell type compositional differences between V1 cell types and wS1 cell types. In a typical workflow, we trained an XGBoost (version 1.3.3) classifier to learn subclass or type labels within the P22 control “reference” dataset and applied this classifier to another “test” dataset. The correspondences between cluster IDs and classifier-assigned labels for the test dataset were used to map subclasses or types between datasets. The classification workflow is described in general terms below and applied to various scenarios throughout the study.

Let *R* denote the reference dataset containing *N_R_*cells grouped into *r* clusters. Let *T* denote the test dataset containing *N_T_* cells grouped into *t* clusters. Each cell is a normalized and log-transformed gene expression vector ***u*** ∈ *R* or ***v*** ∈ *T*. The length of ***u*** or ***v*** equals the number of genes. Based on clustering results, each cell in *R* or *T* is assigned a single cluster label, denoted cluster(***u***) or cluster(***v***). cluster(***u***) may be a type or subclass identity, depending on context.

The main steps are as follows:

1. We trained a multi-class XGBoost classifier *C_R_^0^* on *R* using the intersection of HVGs from *R* and *T* as features. The training dataset was split into training and validation subsets. For training, we randomly sampled 70% of the cells in each cluster, up to a maximum of 1000 cells per cluster. The remaining “held-out” cells were used for evaluating classifier performance. Clusters with fewer than 100 cells in the training set were upsampled via bootstrapping to 100 cells to improve classifier accuracy for the smaller clusters. Classifiers achieved >95% accuracy or higher on the validation set for most clusters, with some clusters yielding 85%-95% accuracy. XGBoost parameters were fixed at the following values:

1. ‘Objective’: ‘multi:softprob’
2. ‘eval_metric’: ‘mlogloss’
3. ‘Num_class’: *r*
4. ‘eta’: 0.2
5. ‘Max_depth’: 6
6. ‘Subsample’: 0.6
2. When applied to a test vector ***c***, the classifier *C_R_^0^* returns a vector *p = (p_1_, p_2_, …)* of length *r*, respectively. Here, *p_i_* represents the probability value of predicted cluster membership within *R,* respectively. These values are used to compute the “softmax” assignment of ***c***, such that cluster(***c***) = *arg max_i_ p_i_* if *arg max_i_ p_i_* > *1.2*(1/r)*. Otherwise ***c*** is classified as ‘Unassigned’.
3. After determining that the initial classifier *C_R_^0^* faithfully learns the reference clusters, we trained a classifier *C_R_* on 100% of the cells in *R*. *C_R_* was then applied to each cell ***v*** ∈ *T* to generate predicted labels cluster(***v***). In this study, *T* was P12, P22 10d AWD, P22 1d RWD, and V1 P21.

The cell type frequency distribution predicted for *T* was compared to the distribution of cell type labels in *R* using scatter plots. Each dot represented one of the clusters in *R,* and the axes represented the frequency of that cluster in each dataset.

#### Principal component analysis on L2/3 pyramidal neurons

The classification analyses revealed that L2/3 was the only subclass where cell type frequency differed significantly between P12 and P22 (**Fig 3B,C** in the main text). To explore this further, we performed principal component analysis on L2/3 cells only. Since we were interested in understanding L2/3 cell type identity and how it varies across conditions, we chose as the features the top markers for each type (Wilcoxon rank-sum test, FC>1.5, FDR<0.05, expressed in >20% of cells in type), resulting in a set of 489 genes. The same set of genes was used for the PCA of all conditions. Correlations between the principal eigenvectors across the conditions were computed by taking their dot product. This is equivalent to the correlation coefficient of the loadings. Note that, by construction, the principal eigenvectors are orthonormal within a sample. Finally, a score for each marker set was computed for each cell as the mean expression of all genes in the set in that cell.

#### Overlap of tDE genes between S1 and V1

To determine the degree to which tDE genes are shared between S1 and V1, we performed subclass-by-subclass differential gene expression on P14 and P21 V1 data, as they are the most closely matched to our P12 and P22 data(10). We then performed a hypergeometric test for the overlap of tDE genes in either direction for each subclass. In each test for each subclass, four variables are set: *N,* the universal set comprising all of the genes downregulated/upregulated in every subclass, *K*, the number of tDE genes in V1, *n*, the number of tDE genes in wS1, and *k*, the intersections of *K* and *n.* The *P* value for each subclass was Bonferroni-corrected by multiplying by the number of subclasses tested.

### Experimental Methods

#### Mice handling

All procedures were approved by the UC Berkeley Animal Care and Use Committee and were in accordance with NIH guidelines. C57Bl6J male mice were obtained from Charles River Laboratories. Mice were maintained on a 12 hr day/night cycle and housed with littermates and the mother in the UC Berkeley animal facility. For whisker deprivation, mice were anesthetized with isoflurane, and whiskers were carefully plucked under a dissecting microscope with forceps using slow and steady force to prevent removal of the whisker follicle. Sham mice were anesthetized for the same amount of time as deprived, but whiskers were only stroked briefly with forceps.

#### Droplet-based snRNA-seq

Mice were anesthetized with isofluorane, rapidly decapitated, and the brain was dissected out into ice-cold Hibernate A (BrainBits Cat# HACA). For each condition (P12, P22 control, P22 1d RWD, P22 10d AWD) 3 mice were used for each biological replicate of single-nucleus(sn) RNA-sequencing. Extracted brains were placed on a metal brain mold (Zivic Instruments,#5569) and the slice containing wS1 was isolated by inserting in the 11th space on the mold (∼7.5 mm from the tip of the olfactory bulb, and a second blade 2 mm further anterior (4 spaces on the mold). This slice was removed and lowered to Hibernate A in a 60cc petri dish, placed on a ruler under a dissecting microscope. The midline was aligned with the ruler, and the first cut was bilaterally 2 mm out from the midline in a radial direction. The second cut was 2 mm medial to the first cut. The cortex was peeled off the underlying white matter. The wS1 piece was transferred into a RNAse-free cryovial, excess liquid was removed, and the tube was rapidly frozen on dry ice. Once all dissections were complete, the tissue was transferred from dry ice into a dewar of liquid nitrogen for storage before nuclei isolation.

#### Nuclei Isolation

Nuclei were isolated using the 10X Chromium nuclei isolation kit (10X Genomics,Cat#1000494). After isolation according to the chromium protocol, cells were counted on a hemocytometer in ethidium bromide and then diluted to 700-1200 nuclei/mm3. Nuclei from each biological replicate were split into two tubes and run separately on two channels of 10X v3, targeting 10,000 cells per channel. We refer to these as library replicates. For each experiment, we performed two or three biological replicates towards a total of four to six library replicates.

#### Fluorescence in situ hybridization (FISH)

C57BL/6 male mice (Charles River), from age P12 and P22, were anesthetized with isoflurane and transcardially perfused with 2% RNAse-free paraformaldehyde (PFA) in PBS (pH 7.4). Mouse brains were collected and immediately fixed in 4% RNAse-free PFA at 4°C. After 24 hours, the brains were transferred to 30% sucrose in PBS and were allowed to sink 4°C. Slices were cut from the left hemisphere in the ‘across-row’ plane. Brains were first mounted on a tissue guillotine with a 35° incline, rostral pointing upward. Brains were then cut at an angle 50° from the midsagittal plane. Using this plane, every S1 slice has one column from each whisker row A–E, and circuitry within columns remains largely intact (38,70). Slices were cut on a sliding microtome (Reichert Scientific Instruments 860) into 30 µm thick sections, collected into RNAse-free PBS, and onto charged microscope slides. Sections were air-dried overnight, then post-fixed in 4% RNAse-free PFA for 1 hour at 4°C; this was followed by serial ethanol dehydration and dried before being promptly stored at −80°C until further processing.

Multiplex fluorescence in situ hybridization followed the protocol for ACDBio’s RNAscope Multiplex Fluorescent V2 Assay (Advanced Cell Diagnostics, cat# 323110). Thawed sections underwent H2O2 permeabilization, 5-minute target retrieval, and protease III treatment. RNAscope probes *Trpc6* (cat# 442951), *Chrm2* (cat# 495311-C2), *Cdh13* (cat# 443251-C3), *Slc17a7* (cat# 501101-C4), *Adamts2* (Cat# 806371-C3)*, Bdnf* (Cat# 424821) *Btg2* (cat# 483001), *Npas4* (cat# 423431-C2), *Junb* (cat# 584761-C3), *Col19a1* (cat# 539701), *Sorcs3* (cat# 473421- C2), *Etv1* (cat# 557891-C3), and *Gabrg3* (cat# 514421-C4), were applied and amplified in sequence with TSA Vivid and Opal Polaris dyes (Advanced Cell Diagnostics, cat# 323271, 323272, 323273; Akoya Biosciences, cat# FP1501001KT). Cellular nuclei were counterstained with 1 µg/ml DAPI and mounted with Prolong Glass Antifade Mountant (Thermo Fisher Scientific, cat# P36982). All RNAscope runs were performed with both conditions side-by-side and controls, to reduce variability.

Imaging was performed using a Zeiss Axio Scan 7 slide scanner with Zen digital imaging software. Tilescan images were acquired of the entire wS1 including all cortical layers at 20X. All channels were acquired at the same exposure, gain and laser power settings for each condition. The relevant barrel columns were identified using DAPI staining in combination with the other markers and images were cropped to target L2/3 and barrel columns of interest in FIJI. All pre-processing steps were kept consistent between conditions. Cropped images were inputted into CellProfiler to detect 5 imaging channels. Nuclear segmentation was performed with DAPI channel and cellular segmentation was performed with the RNAscope probe *Slc17a7* (vGlut1), a marker for glutamatergic neurons. For analysis of temporally-regulated mRNAs, nuclear segmentation with DAPI was used as a region of interest (ROI) to measure mean intensity values for each mRNA in each nucleus. Layers were estimated using empirically measured distance from the pial surface. L2/3 was considered 50-250 µm deep, and L4 was 250-350 µm. For cell type analysis, thresholds were set to identify cells expressing markers above background. Threshold parameters for each channel were kept the same across conditions. To determine cell type identity and overlap, cell type markers were identified as objects. Objects that did not overlap with the cell segmentation marker *Slc17a7* were eliminated in order to isolate excitatory L2/3 cells. Lastly, object overlap across channels was computed. The coordinates of all identified objects from CellProfiler were used to generate scatterplots in Python with the Seaborn package. For immediate-early genes (IEG) analysis, cellular segmentation with *Slc17a7* was used as a region of interest (ROI) to measure intensity values for each mRNA in each excitatory L2/3 cell. Outliers were cleaned from the data in Graphpad Prism using the ROUT method(71). Outline plots were generated in CellProfiler(72), by relating cellular segmentation object results and IEG objects. Outlines for each image used in the analysis were overlaid by z-projecting in FIJI to generate summary outline figures. Plots and statistics were generated using Graphpad Prism 10. Normality tests were performed on all datasets. If datum were not normally distributed, nonparametric statistical tests were used. Each condition (P12, P22 control, P22 1d sham, P22 1d RWD) had 3-5 mice from which multiple brain sections were analyzed.

### Data and code availability

All raw and processed snRNA-seq datasets reported in this study will be made publicly available via NCBI’s Gene Expression Omnibus (GEO) Accession Number GSE276528. Processed h5ad files are available at https://github.com/shekharlab/wS1dev. These h5ad files contain all relevant metadata and log-normalized counts. Computational scripts detailing snRNA-seq analysis reported in this paper are available at https://github.com/shekharlab/wS1dev. All custom software for imaging analysis will be made available upon request. Any additional information required to reanalyze the data reported in this paper is available from the corresponding authors upon request.

## Acknowledgments

Select images were made using BioRender. We thank Dr. Justin Choi from the QB3/Functional Genomics Lab (RRID:SCR_022170) for assistance with scRNA-seq. We also thank Drs. Melanie Oakes and Quy Nguyen from the UCI sequencing core for performing all sequencing for this study. The UCI Genomics Research and Technology Hub (GRT Hub) is partly supported by NIH grants to the Comprehensive Cancer Center (P30CA-062203) and the UCI Skin Biology Resource Based Center (P30AR075047) at the University of California, Irvine, as well as to the GRT Hub for instrumentation (1S10OD010794-01and 1S10OD021718-01). We are grateful to Drs. Juyoun Yoo and Saumya Jain from the UCLA Zipursky lab for feedback during the early stages of nuclei extraction protocol optimization. This work was funded by NIH grants 1F32 NS126310 (HRM), 1F31 F31NS131016 (SB), EY028625 (KS), R01 NS105333 (DF), as well as funds from the Society of Hellman Fellows (KS), the McKnight foundation (KS), and SFARI Investigator Award grant (DF).

## Author Roles

S. B., Conceptualization, Data curation, Formal analysis, Investigation, Methodology, Validation, Visualization, Software, Writing – original draft, Writing – review & editing.

H.M., Conceptualization, Data curation, Formal analysis, Investigation, Methodology, Validation, Visualization, Writing – review & editing

C.Y., Investigation

D.F., Conceptualization, Methodology, Validation, Writing – review & editing, Resources, Funding Acquisition, Supervision

K.S., Conceptualization, Methodology, Validation, Writing – review & editing, Resources, Funding Acquisition, Supervision

S.B. and H.M. contributed equally to the study.

## SUPPORTING INFORMATION CAPTIONS

**Figure S1.**
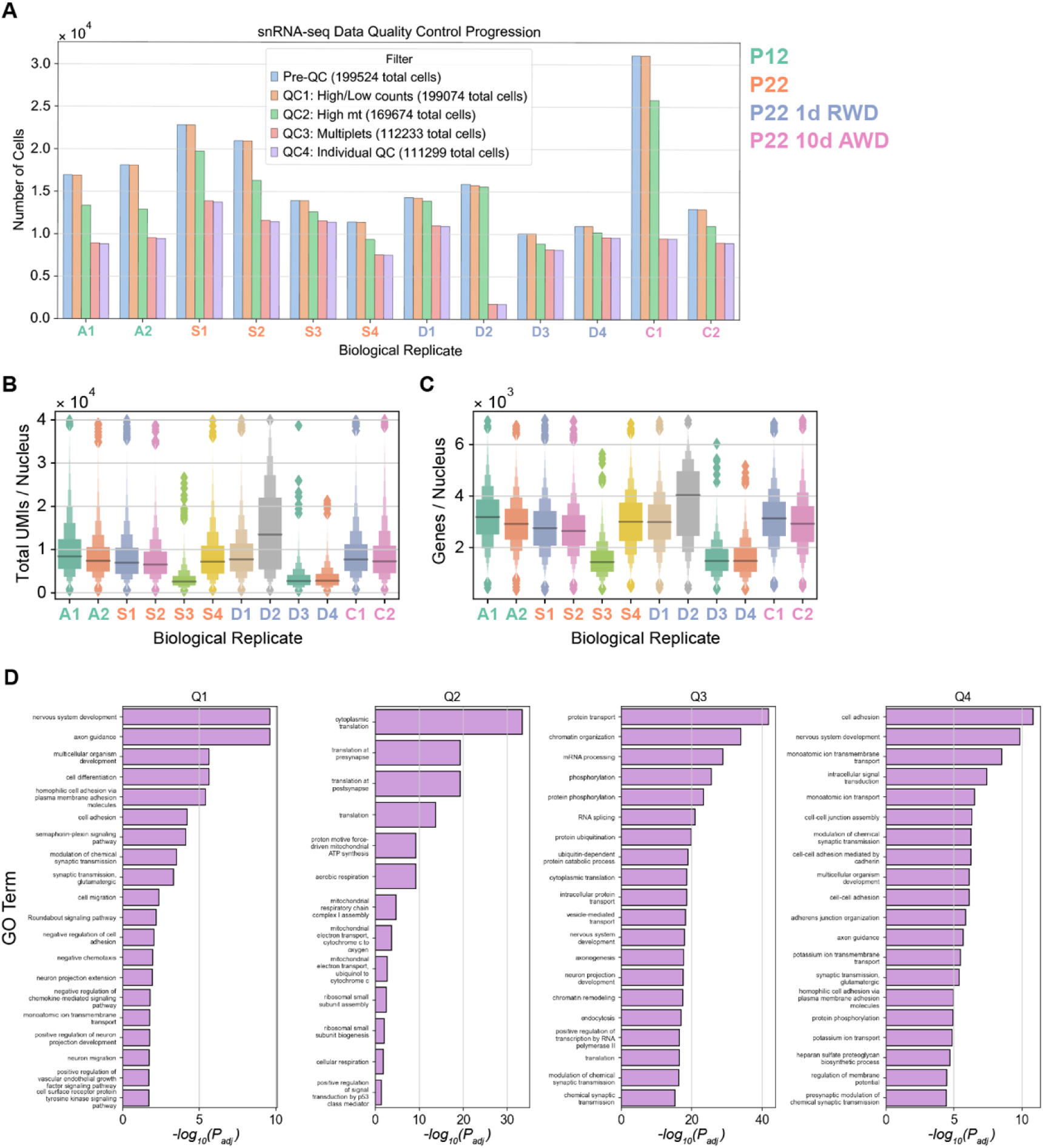
Data filtering steps, quality control, and gene ontology (GO) analysis of temporally regulated genes. A. Bar plots showing the number of nuclei remaining in each biological replicate at the end of each filtering step (see **Methods** for details). Biological replicates (x-axis) are colored by their experimental condition (legend, right). “PreQC” represents the default number of nuclei the 10X CellRanger software provides. “QC4” represents the final set of nuclei used for downstream analyses. B. Distribution of total RNA counts detected in each biological replicate from each condition. C. Distribution of total number of genes detected in each biological replicate from each condition. D. The top 20 “biological process” gene ontology terms for Q1-Q4 for glutamatergic subclasses as shown in **Fig 2D**.

**Figure S2.**
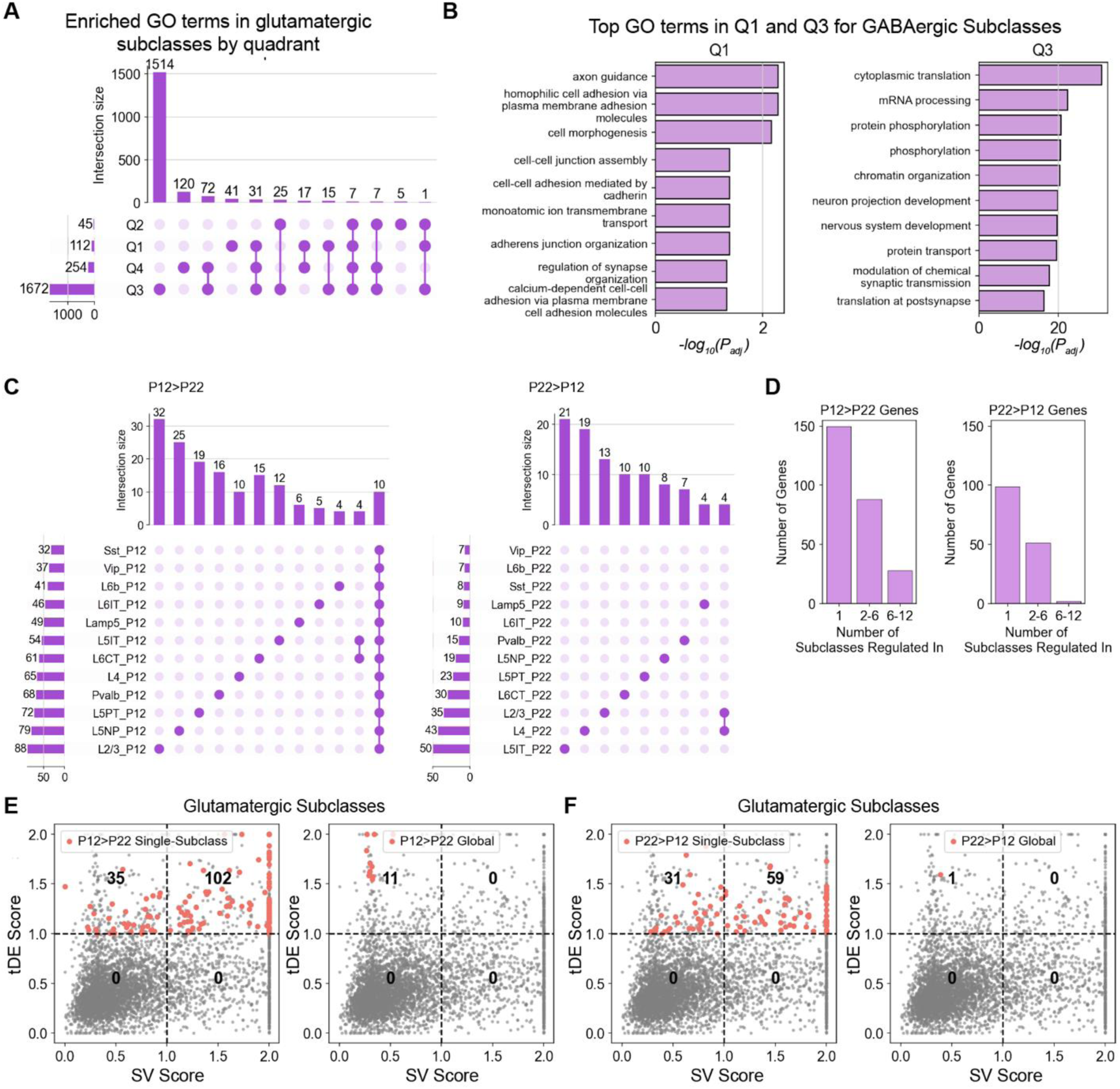
Developmental gene expression changes are subclass-specific. A. UpSet plot (69) showing the overlap of GO terms associated with “biological process” (BP) across Q1-Q4 for glutamatergic neuronal subclasses. The lower panel indicates the set intersections corresponding to each column (e.g. the third column indicates the number of GO terms found in Q3 and Q4, but not in Q1 and Q2). B. Top GO terms enriched in Q1 (*left*) and Q3 (*right*) for GABAergic neuronal subclasses. C. UpSet plots showing that downregulated (*left*) and upregulated (*right*) tDE genes between P12 and P22 are primarily subclass-specific. Only set intersections containing at least four genes are shown. Note that, unlike panel A, the sets here correspond to groups of subclasses rather than groups of quadrants. D. Bar plots summarizing that ∼60% of genes are regulated in only one subclass and that the number of downregulated genes is ∼1.6x that of upregulated genes. E. Visualization of the subclass-specific and global P12>P22 genes from panel A in the quadrant analysis of **Fig 2B** for glutamatergic neurons. F. Same as panel E for P22>P12 genes.

**Figure S3.**
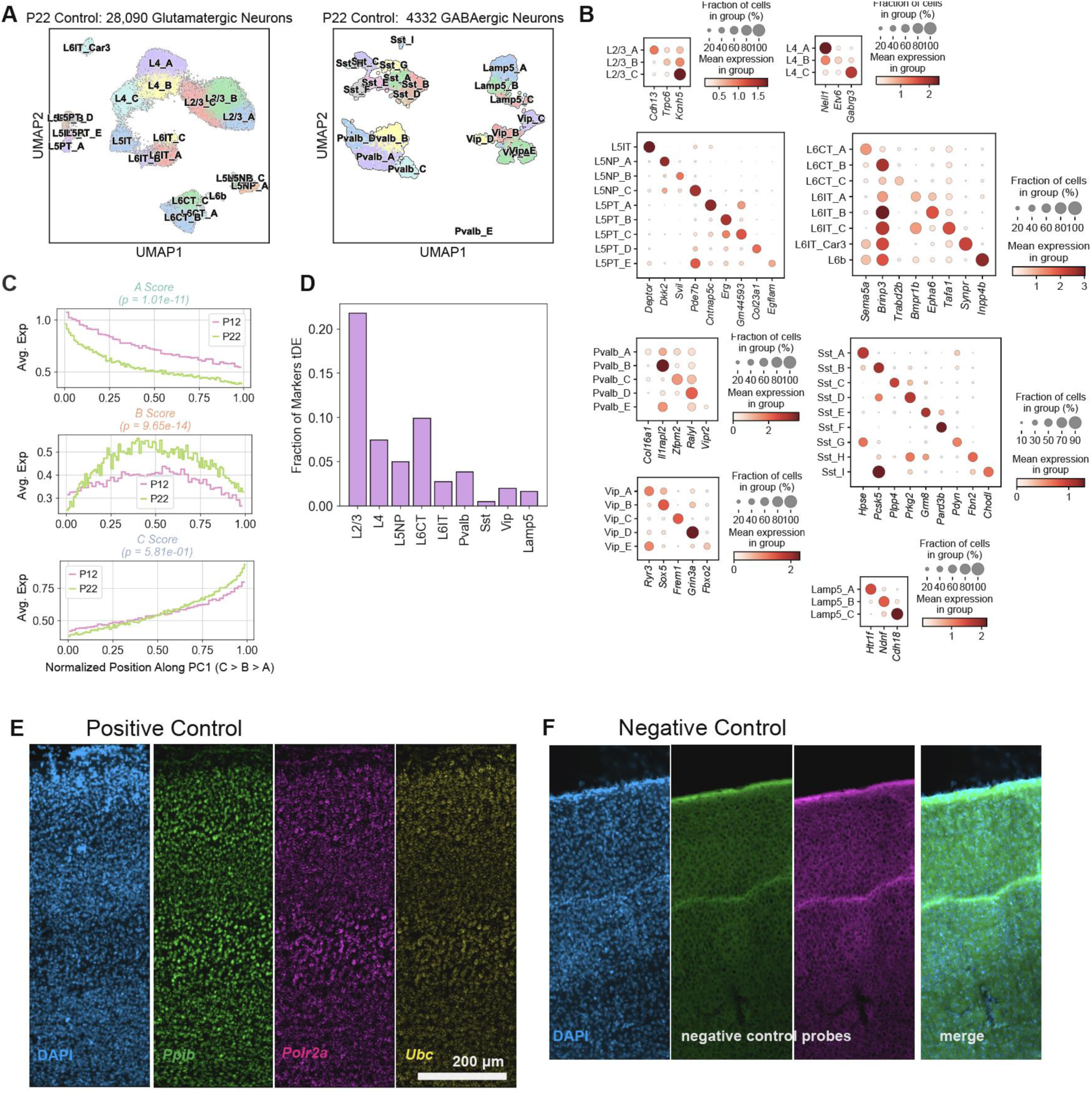
Neuronal cell types at P22 and developmental changes. A. UMAP visualization of P22 wS1 cell types in glutamatergic (left) and GABAergic (right) neurons. B. Dotplots showing top cell type markers within each subclass at P22. Within each dotplot panel, rows indicate cell types and columns indicate genes. The size of each circle corresponds to the % of cells with nonzero expression, and the color indicates average expression level. C. Same as **Fig 4A**, but the y-axis now plots aggregate expression scores for_L2/3_A, L2/3_B, and L2/3_C along PC1. Curves correspond to P12 and P22. *P*-values are from a Kolmogorov–Smirnov test between the two ages. D. Barplot showing that L2/3 has more markers that are tDE between P12 and P22 than the other subclasses. E. Representative widefield images of RNAscope positive control showing expected labeling pattern. F. Representative widefield images of RNAscope negative control using nontargeting probes showing no signal as expected.

**Figure S4.**
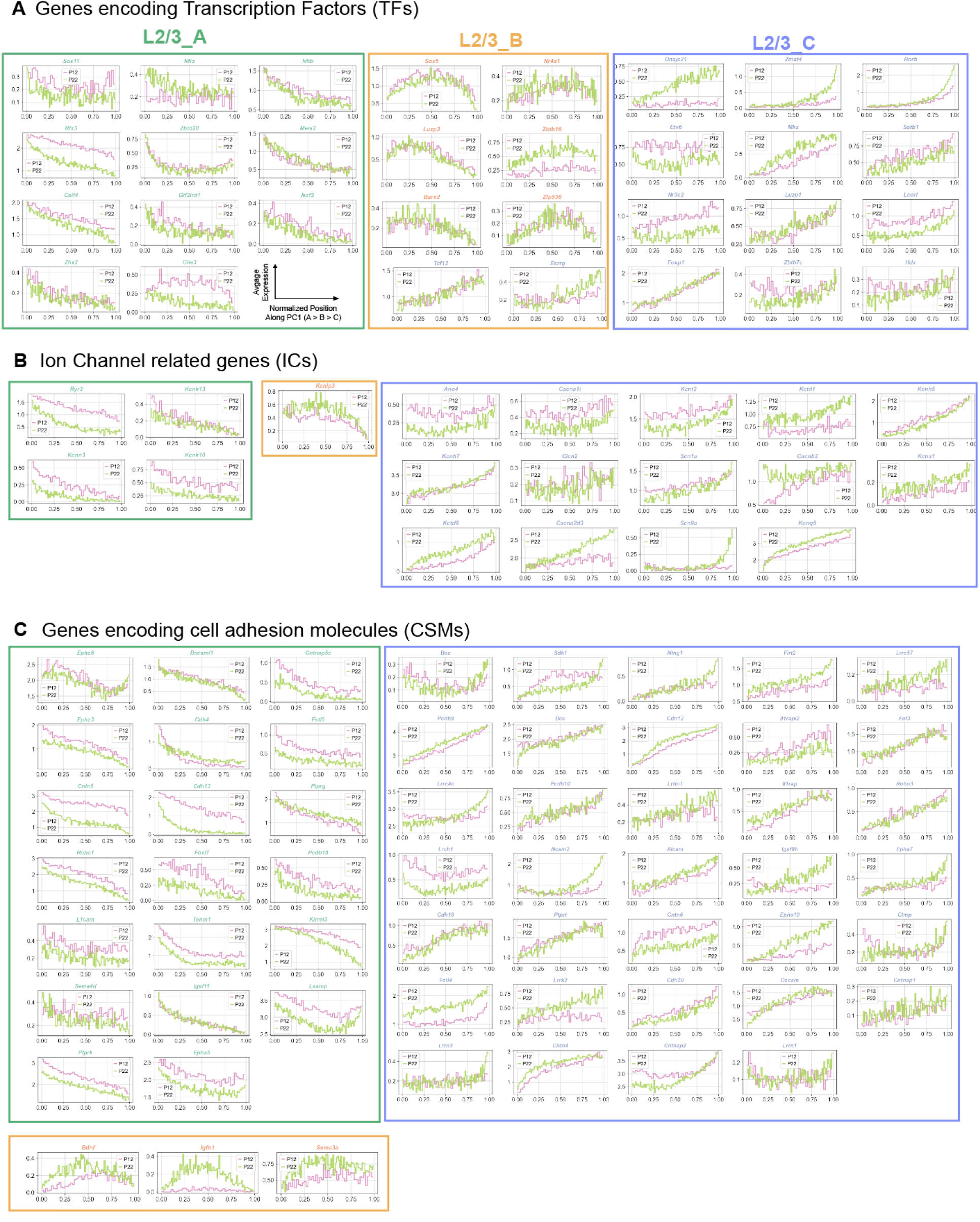
Expression patterns at P12 and P22 along PC1 of L2/3 type-enriched genes related to transcription factors (TFs), cell adhesion molecules (CAMs), and ion channels (ICs). A. Expression patterns of type-enriched TFs at P12 and P22 in L2/3 cells ordered by PC1 value. B. Same as A for ICs C. Same as A for CAMs

**Figure S5.**
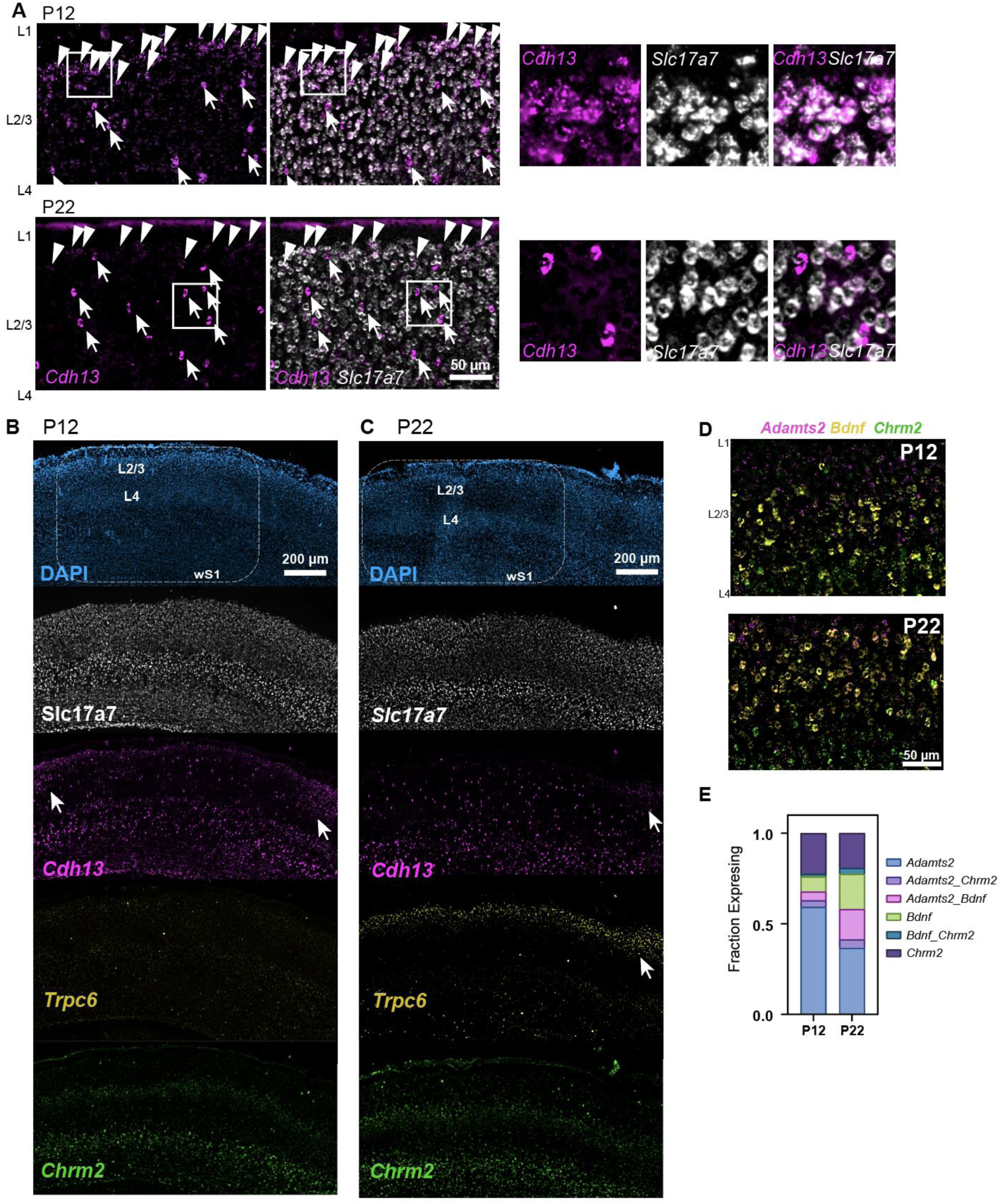
Representative FISH images of L2/3 cell type markers. A. Representative images of *Cdh13* labeling at P12 (top row) and P22 (bottom row). Overlay with *Slc17a7* (vGlut1) shows that the majority of *Cdh13* expressing cells in the middle of L2/3 do not colocalize with *Slc17a7* (white arrows) whereas the *Cdh13+* cells along the Layer 1/2 border do coexpress *Slc17a7* (white arrowheads). (right) Inset from area inside white squares. B. Widefield images of ‘across-row’ section (see Methods for details) with wS1 and surrounding cortical areas at P12. Arrows indicate cortical regions outside of wS1 where labeling becomes denser. C. Widefield images of ‘across-row’ section with wS1 and surrounding cortical areas at P22. Arrows indicate cortical regions outside of S1 where labeling becomes denser. D. Representative FISH images of wS1 L2/3 labeling cell type markers *Adamts2, Bdnf,* and *Chrm2* at P12 and P22. E. Quantification of the fraction of excitatory (*Slc17a7+*, not shown) L2/3 cells expressing one or more of markers *Adamts2*, *Bdnf*, and *Chrm2* at P12 and P22. N = 3-4 slices from 2 mice per time point.

**Figure S6.**
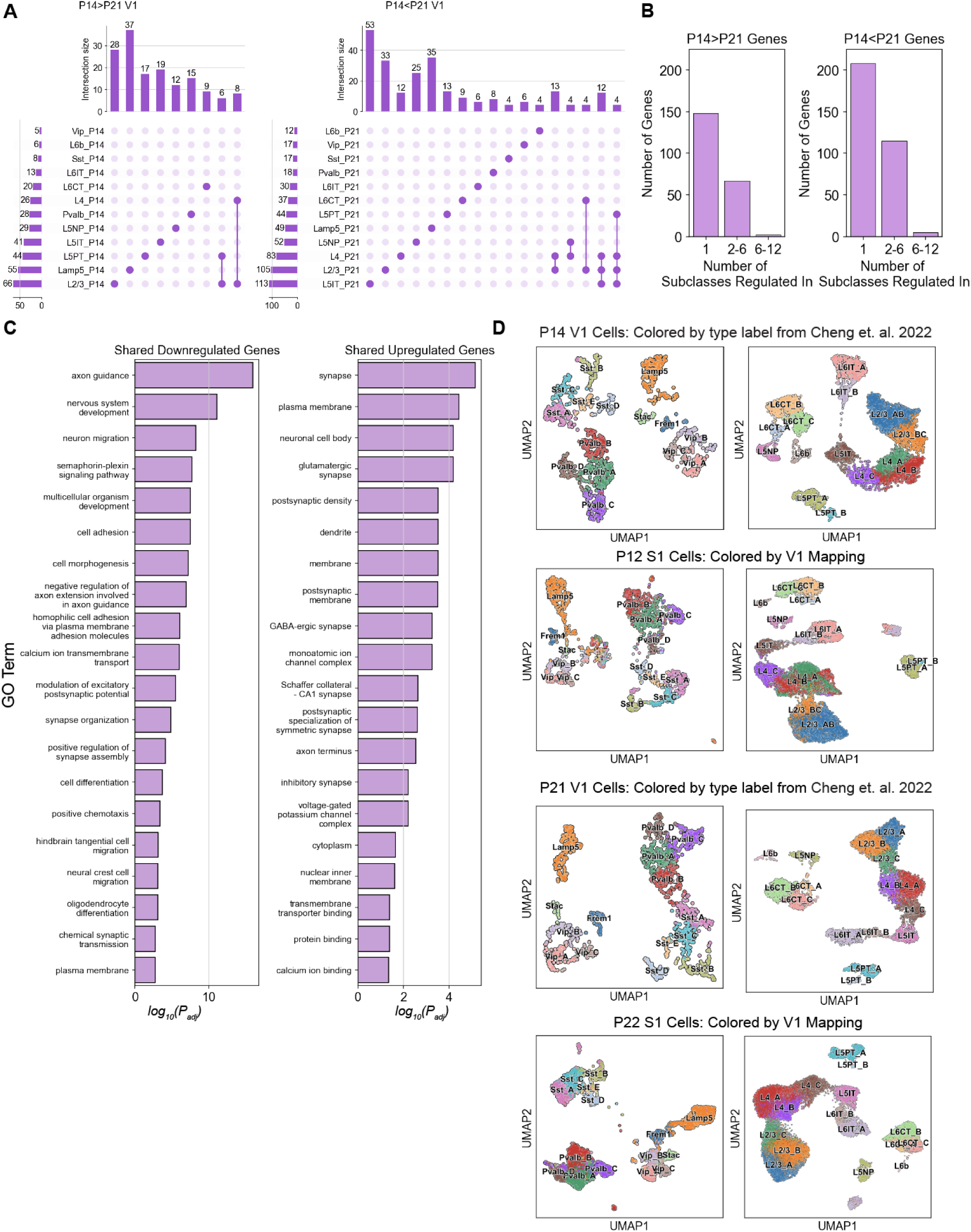
Temporal gene expression changes in V1, GO enrichment for shared and region-specific genes, and mapping analysis. A. UpSet plot (as in **Fig S2C**) summarizing subclass-by-subclass tDE analysis of V1 data. Only combinations containing at least four genes are shown. B. Barplots showing that as in the case of wS1 (**Fig S2D**), most downregulated (*left*) and upregulated (*right*) genes in V1 are subclass-specific. C. Full list of GO terms enriched in shared downregulated (*left*) or upregulated (*right*) tDE genes between V1 and wS1. D. UMAP plots of V1 (rows 1 and 3) and wS1 (rows 2 and 4) data colored by V1 labels. V1 neuron labels are based on the published clustering in Cheng et al. (10), while wS1 neurons were labeled using a supervised mapping analysis (see **Methods**).

**Figure S7.**
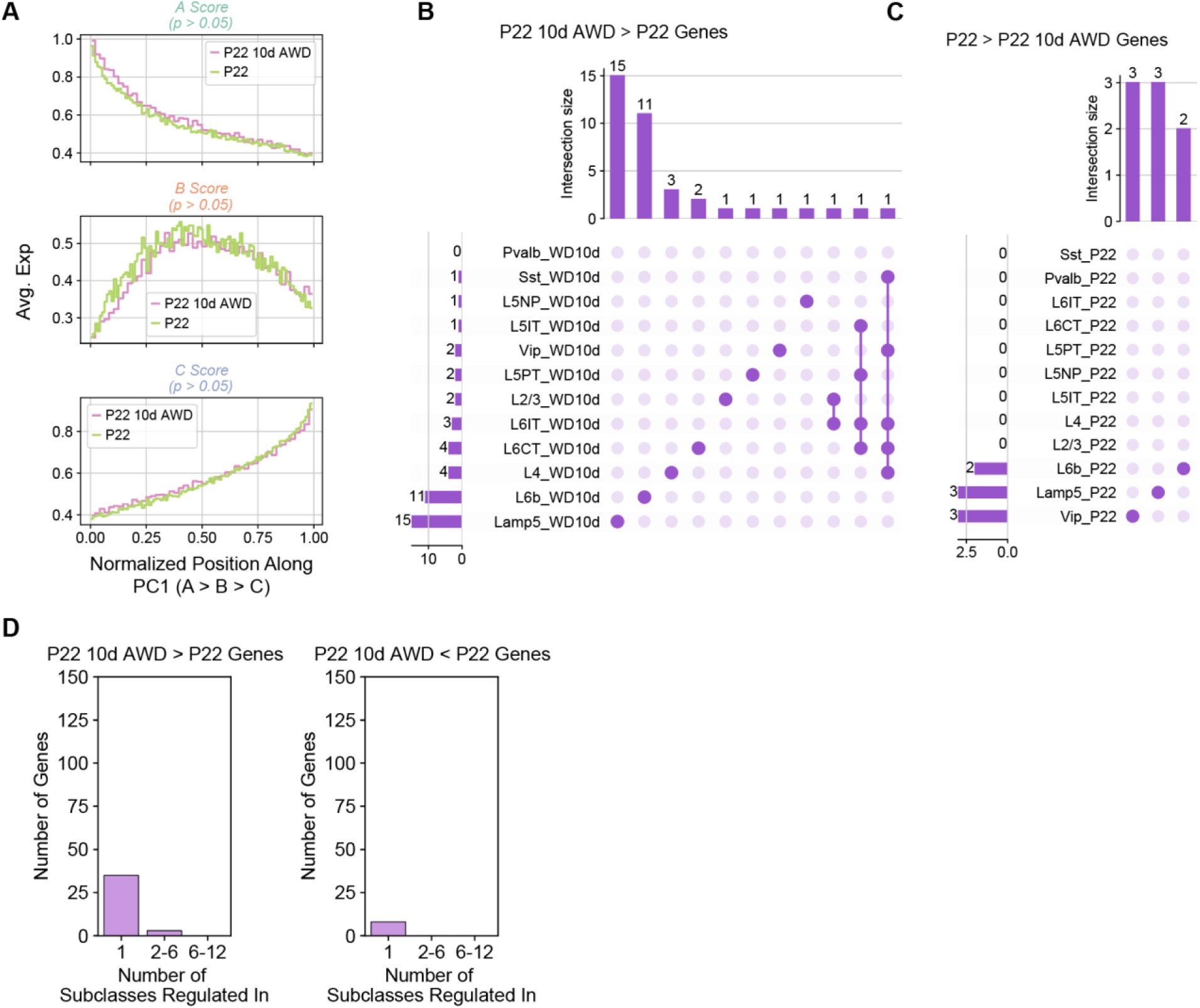
Subclass-level gene expression changes between P22 10d AWD and P22 control. A. L2/3 type A, B, and C marker scores plotted as a function of a cell’s position along PC1. *P* values are based on a Kolmogorov–Smirnov test comparing the two conditions. B. UpSet plots showing that the few genes upregulated by 10d AWD are predominantly subclass-specific. C. Same as B but for genes downregulated by 10d AWD. D. Bar plots highlighting the small number of genes regulated by 10d AWD. Scale for y-axis is the same as for **Fig S2D** for comparison.

**Figure S8.**
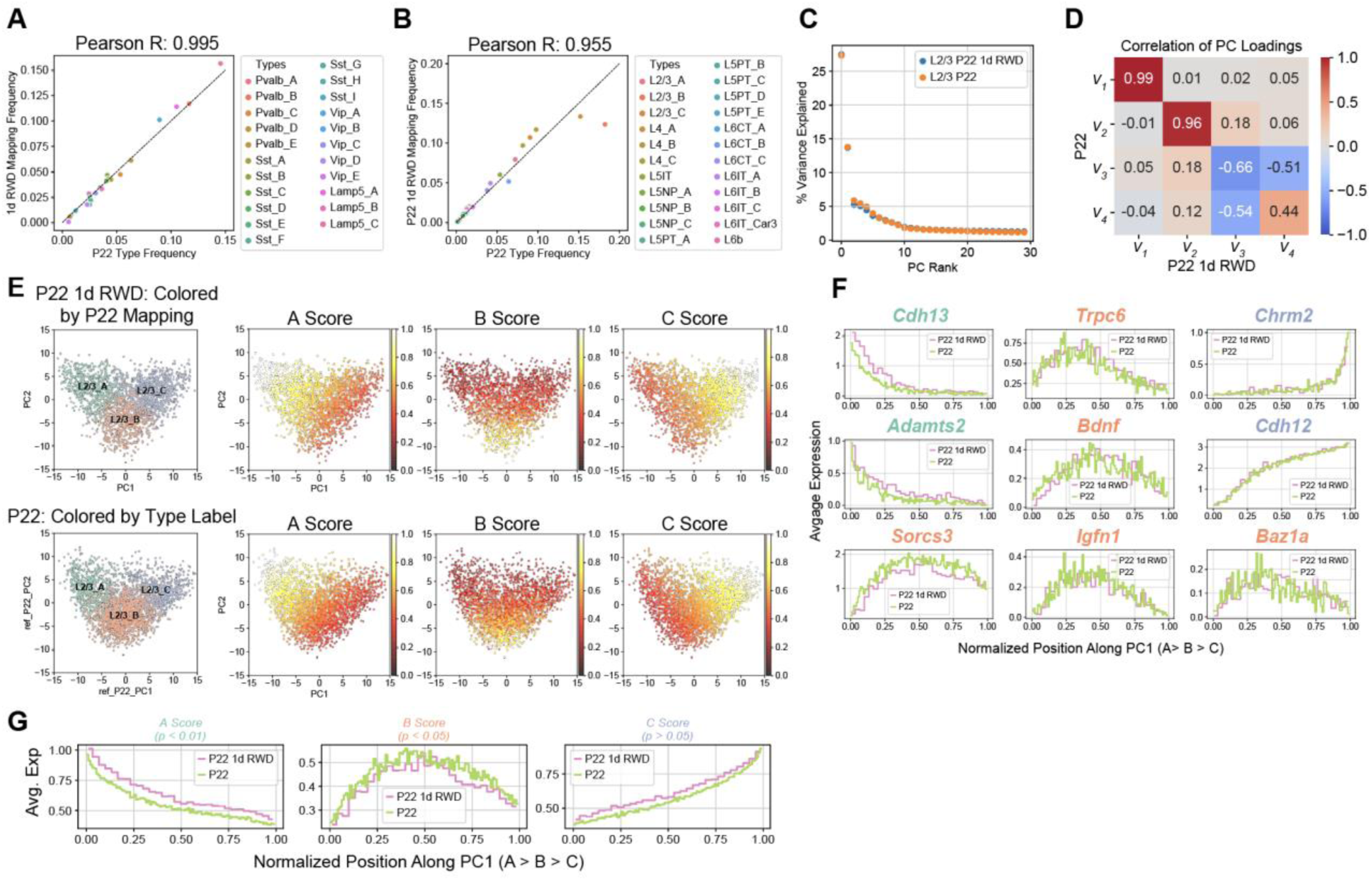
1d RWD has little effect on L2/3 cell type identity. A. GABAergic cell types have approximately the same relative frequency at P22 1d RWD and P22 normal whisker experience. Representation as in **Fig 6A**. B. Same as panel A, for glutamatergic cell types. C. PC1 and PC2 are sufficient to describe transcriptional variance within L2/3 in the normal P22 and P22 10d AWD datasets D. Similar to **Fig 6D**, comparing principal eigenvectors between the P22 1d RWD and normal P22 datasets. The first two principal eigenvectors map 1:1. E. Similar to **Fig 6E** comparing the PC1 vs. PC2 distribution and type-specific scores between P22 1d RWD and normal P22 L2/3 datasets. F. L2/3 markers genes, as in **Fig 4A**, are shown as a function of cells’ position along PC1 comparing patterns between normal P22 and P22 1d RWD. G. L2/3 type A, B, and C marker scores plotted as a function of a cell’s position along PC1. *P* values are from a Kolmogorov– Smirnov test between the two conditions.

## Table Legends

**Table S1.** tDE and SV scores for every tested gene in glutamatergic and GABAergic neurons.

**Table S2.** Subclass-by-subclass differential expression testing results between P12 and P22.

**Table S3.** Cell type markers from each subclass at P22.

**Table S4.** Subclass-by-subclass differential expression testing results between P14 and P21 V1 data.

**Table S5.** Subclass-by-subclass differential expression testing results between P22 10d AWD and P22

**Table S6.** Subclass-by-subclass differential expression testing results between P22 1d RWD and P22

